# Quantitative imaging of transcription in living *Drosophila* embryos reveals the impact of core promoter motifs on promoter state dynamics

**DOI:** 10.1101/2021.01.22.427786

**Authors:** Virginia L Pimmett, Matthieu Dejean, Carola Fernandez, Antonio Trullo, Edouard Bertrand, Ovidiu Radulescu, Mounia Lagha

## Abstract

Genes are expressed in stochastic transcriptional bursts linked to alternating active and inactive promoter states. A major challenge in transcription is understanding how promoter composition dictates bursting, particularly in multicellular organisms. We investigate two key *Drosophila* developmental promoter motifs, the TATA box (TATA) and the Initiator (INR). Using live imaging in *Drosophila* embryos and new computational methods, we demonstrate that bursting occurs on multiple timescales ranging from seconds to minutes. TATA-containing promoters and INR-containing promoters exhibit distinct dynamics, with one or two separate rate-limiting steps respectively. A TATA box is associated with long active states, high rates of polymerase initiation, and short-lived, infrequent inactive states. In contrast, the INR motif leads to two inactive states, one of which relates to promoter-proximal polymerase pausing. Surprisingly, the model suggests pausing is not obligatory, but occurs stochastically for a subset of polymerases. Overall, our results provide a rationale for promoter switching during zygotic genome activation.

## Introduction

In all eukaryotes, transcription of active genes into RNA requires the controlled assembly of multiple protein complexes at promoters^1^. This includes the sequential recruitment of general transcription factors to form the pre-initiation complex (PIC), followed by the recruitment of RNA Polymerase II (Pol II) at the transcriptional start site (TSS)^2–4^ of promoters. Among all these complexes, TFIID stands out as the primary core promoter recognition factor that triggers PIC assembly^5^. TFIID includes TATA binding protein (TBP), which binds upstream of the TSS, as well as 13 TBP-associated factors (TAFs), known to bind downstream promoter elements such as the Initiator element (INR) and the Downstream Promoter Element (DPE) motifs^3,6^.

After initiation, Pol II transcribes a short stretch of 30-80 nucleotides before pausing^7^. This step is regulated by the TFIID complex, as directly demonstrated *in vitro*^8^ and inferred from the over-representation of TFIID-bound core promoter motifs (INR, DPE, pause button) in highly paused genes^9–13^. Pausing durations are highly variable among genes^14–16^. However, it is thought that exit from the paused state is a kinetic bottleneck in the transcriptional cycle^17^ and is often used as a checkpoint during development to foster coordination in gene activation, plasticity or priming^18–21^. How core promoter motifs affect this rate-limiting step is still unknown, particularly in the context of a developing embryo.

In parallel to these genomics approaches, single cell imaging revealed that transcription is not a continuous process over time but occurs through stochastic fluctuations between periods of transcriptional activity and periods of inactivity, called bursting^22^. Direct labeling of newly synthesized RNA with the MS2/MCP amplification system^23,24^ has been the method of choice to observe these bursts in real time in living cells^25–27^ or multicellular organisms^28,29^. In the context of a developing embryo, these studies revealed how enhancers regulate bursting during pattern formation^30–32^. However, relatively less attention has been given to the impact of core promoter motifs on transcriptional bursting.

To build a mechanistic understanding of bursting beyond a qualitative description of burst size and frequency, it is critical to consider the various timescales of the transcription process and to employ mathematical modeling explicitly describing various promoter states^22^. Indeed, depending on the gene and the cellular context, transcriptional bursting has been shown to occur at multiple timescales^33,34^, from seconds (e.g polymerase clusters, Pol II firing^35–37^), to minutes (e.g TBP binding/unbinding, transcription factor binding^38,39^) to hours (e.g nucleosome remodelling, chromatin marks^37,40^). These different timescales can all occur at a single gene, and the term multiscale bursting was coined to describe such complex bursting kinetics^34,37^. This multiscale bursting might explain why the simple two state model (random telegraph), whereby promoters switch from an active to an inactive state, does not always suffice to reliably capture promoter dynamics^36,37,41,42^.

While it is well understood that *cis*-regulatory sequences must impact transcriptional bursting, with enhancers primarily affecting burst frequencies^43,44^, a detailed dissection of the impact of promoter motifs on promoter state dynamics is lacking. In this study we sought to examine the role of core promoter sequences on transcriptional bursting in *Drosophila* embryos. In particular, we focus on two core promoter motifs, the TATA box (TATAWAWR) and the INR (TCAGTY in *Drosophila*^6^) which represent pivotal core promoter contact points with the TFIID complex^45,46^, and are known to regulate both initiation and promoter pausing^8,12^.

The *Drosophila* early embryo is an ideal system to decipher transcriptional bursting regulation since spatially distinct patterns of gene expression are deployed within a relatively short time frame in a multinucleated syncytium, rapidly dividing, that is highly amenable to quantitative imaging^47^. At this early stage of development, the zygotic genome, initially transcriptionally silent, awakens progressively while cell cycle durations lengthen, a process known as the maternal-to-zygotic genome transition (MZT)^48^. This zygotic genome activation (ZGA) occurs gradually through two waves, a major wave during nuclear cycle 14 (nc14) and a minor wave occurring at earlier stages. Moreover, *Drosophila* developmental genes show a clear promoter code with well defined promoter elements^6,49^, with differential usage during ZGA^50^.

We employed the MS2/MCP system to monitor nascent transcription. We implemented a machine learning method that deconvolves single nuclei mRNA production from live *Drosophila* embryos to detect all single polymerase initiation events. The waiting times between successive initiation events were further analysed to infer the number of promoter states and the transition rates among these states. Our results show that TATA box-containing promoters are highly permissive to transcription through long ON durations and high Pol II firing rates. However, the presence of an INR in the core promoter necessitates the use of a three-state model, with a second inactive promoter state that is likely a consequence of Pol II pausing. We propose a renewed view of promoter pausing whereby only subsets of polymerases enter into a paused state.

## Results

### A synthetic platform to image promoter dynamics in living embryos

To examine how the core promoter sequence influences gene expression variation, we developed a synthetic platform whereby several core promoters were isolated, cloned into a minigene and inserted at the same site in the *Drosophila* genome (**Figure 1A**). This approach allows for a direct comparison of core promoter activity since differences caused by variations in *cis*-regulatory context, genomic positioning and mRNA sequence (particularly the 3’UTR) are eliminated. Core promoters were inserted immediately downstream of the *snail* (*sna*) distal minimal enhancer (*snaE*)^51^. The core promoters were selected based on the presence of known core promoter motifs^49^ such as a TATA box in *sna* or an INR in *kruppel* (*kr*) and *Insulin-like peptide 4* (*Ilp4*) (**Figure 1A**, **Supplemental Figure 1, Supplemental Movies S1** and **S5**). We also selected the *brinker* (*brk*) core promoter, which is devoid of any known canonical promoter motifs. None of these four promoters have a DPE, however *Ilp4* possesses a bi-partite bridge element^49^. These developmental genes are endogenously expressed during nc14^52,53^ and precise TSS positions were established using embryonic CAGE (Cap Analysis of Gene Expression) datasets^54^ (**Supplemental Figure 1)**. For all transgenes, we used 100bp of promoter sequence, as previous work established that minimal promoter sequences are sufficient to establish pausing *in vivo*^12,21^.

**Figure 1:**
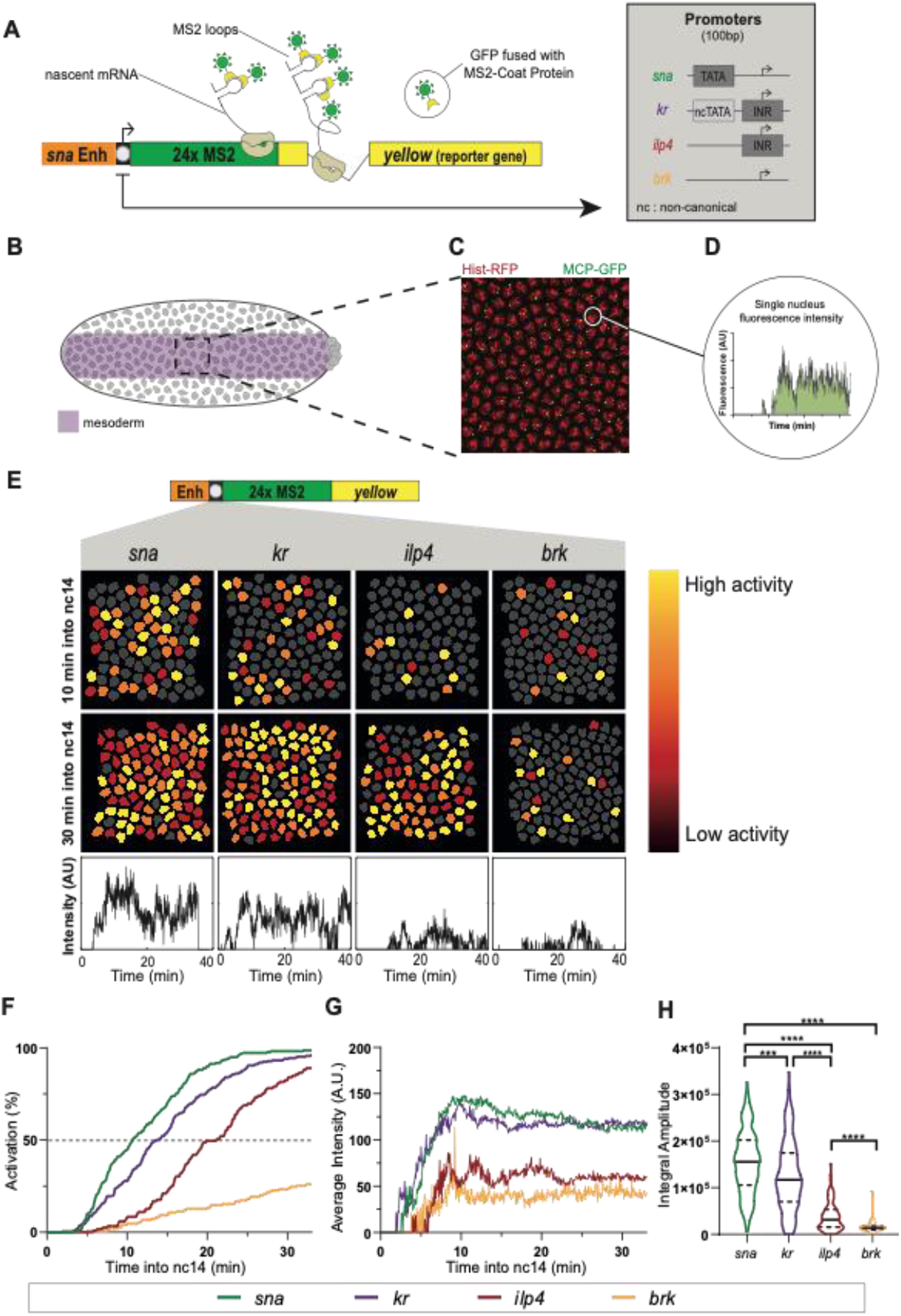
A synthetic transgenic platform to image promoter dynamics. **A)** Schematic view of transgenes used to study transcriptional dynamics of *sna*, *kr*, *Ilp4* and *brk* core promoters. A minimal *sna* enhancer was placed upstream of the core promoter followed by 24xMS2 repeats and a *yellow* reporter gene. Core promoter motifs are indicated in inset. **B)** Schematic of *Drosophila* embryo showing spatial restriction of analysis to presumptive mesoderm (purple). **C)** Maximum intensity projection of representative 15μm Z-stack of *snaE<snaPr<24xMS2-y* (*snaE<sna*) nc14 embryo showing MS2/MCP-GFP-bound transcriptional foci (GFP) and nuclei (histone-RFP). Scale bar is 5μm. **D)** Sample single nuclei trace showing GFP fluorescence during nc14. Surface of green region indicates trace integral amplitude. **E)** False-coloured frames from live imaging of indicated promoters showing relative instantaneous fluorescence intensity in early and late nc14. Inactive nuclei are grey. **F)** Cumulative activation curves of all nuclei during the first 30 minutes of nc14. Time zero is from anaphase during nc13-nc14 mitosis. **G)** Average instantaneous fluorescence of transcriptional foci of active nuclei during the first 30 minutes of nc14. Time zero is from anaphase during nc13-nc14 mitosis. **H)** Distribution of individual trace integral amplitudes from first 30 minutes of nc14. The intensity amplitude at a given time may result from the overlap of several bursts and is a convolution of promoter active/inactive times, polymerase initiation frequency, and the duration of a single polymerase signal. The integral amplitude estimates the transcriptional activity and the total number of transcripts at steady state; it is proportional to the probability of active state (pON) initiation rate (k_INI_) and to the duration of the signal. Solid lines represent median and dashed lines first and third quartiles, using a Kruskal-Wallis test for significance with multiple comparison adjustment. *** p = 0.0002, **** p < 0.0001. **Statistics:** *snaE<sna*, 216 nuclei, 3 movies; *snaE<kr* 243 nuclei, 4 movies; *snaE<ilp4*, 114 nuclei, 2 movies; *snaE<brk*, 45 nuclei, 2 movies. See **Supplemental Movies S1 and S5**.

To track transcription, 24 MS2 stem loops were placed in the 5’ UTR downstream of the promoter, followed by the sequence of the *Drosophila yellow* gene, a gene fragment used as a reporter because of its large size and lack of endogenous expression in the early embryo^55^. MS2 stem-loops in transcribed mRNA were bound by a maternally-supplied MCP-GFP fusion protein, which allowed the detection of transcriptional foci as bright GFP spots. We imaged living *Drosophila* embryos at a high temporal resolution of one frame per 3.86 seconds and focused on the first 30 minutes of nc14^56^. To ensure that all quantified nuclei experience similar peak levels of transcriptional activators, we restricted our analysis to a defined domain within the mesoderm (25μm either side of the presumptive ventral furrow; **Figure 1B**)^57^. Signal intensity at the transcription site was retrieved for each nucleus after 3D detection and tracked throughout nc14 (with mitosis between nc13 and nc14 considered as time zero; see Methods section) and then associated with their nearest nucleus. This generated individual 4D nuclear trajectories (**Supplemental Figure 2**; **Methods**).

### Core promoters differentially affect transcriptional synchrony

We hypothesized that differences in core promoter motifs would result in variability in gene expression independently of specific enhancer contacts. To test this, we characterized single nucleus transcriptional activity (**Figure 1C-E**), as well as the initial timing of activation. As previously reported with fixed sample approaches^21,58^, we observed a spectrum of synchrony profiles (**Figure 1F**), defined as the temporal coordination of activation among a spatially defined domain. The TATA-driven *sna* and INR-driven *kr* minimal promoters showed a rapid activation with a t50 (the time needed for 50% of nuclei within the region of interest to show GFP puncta) of 11 minutes and 14 minutes, respectively while the INR-driven *Ilp4* promoter was delayed with t50 of 21 minutes. The *brk* core promoter did not surpass 24% of nuclei activated throughout nc14.

In addition to differences in synchrony, the overall mRNA production also varied between transgenes (**Figure 1G**). Within our four developmental promoters, promoters showing more rapid reactivation in nc14 also had higher TS intensity in active nuclei, indicating increased instantaneous mRNA production. To examine the total mRNA output, we looked to the integral amplitude, or the area of the curve defined by the average MCP-GFP signal plotted over time (**Figure 1D**). The integral amplitude showed that promoters mediating faster activation and higher instantaneous mRNA production had a higher total mRNA output (**Figure 1H**).

Thus, we established an experimental set-up and quantification pipeline that allows disentangling the contribution of minimal promoter sequences on transcriptional activation of individual, naturally synchronized nuclei within a well-defined, homogeneous spatial domain in live embryos. This initial quantitative comparison of four natural developmental promoters suggests that those with a canonical promoter motif (either TATA or INR) tend to produce higher levels of expression.

### A machine learning method to infer promoter state transition rates

With current labeling and imaging technology, individual transcriptional initiation events cannot be easily and directly visualized, tracked and quantified in the early *Drosophila* embryo. Live imaging of transcription typically shows spots comprised of several newly synthesized mRNAs resulting from the action of multiple polymerases. In order to calibrate fluorescent signal from live imaging, we used single-molecule hybridization experiments^28^. Using the fluorescence of a single mRNA molecule, we estimated the average number of mRNA molecules present at the TS within the nucleus at steady state (**Supplemental Figure 3**). This calibration step allowed us to express single nuclei TS intensities as an absolute number of transcribing polymerases (**Figure 2A**).

**Figure 2:**
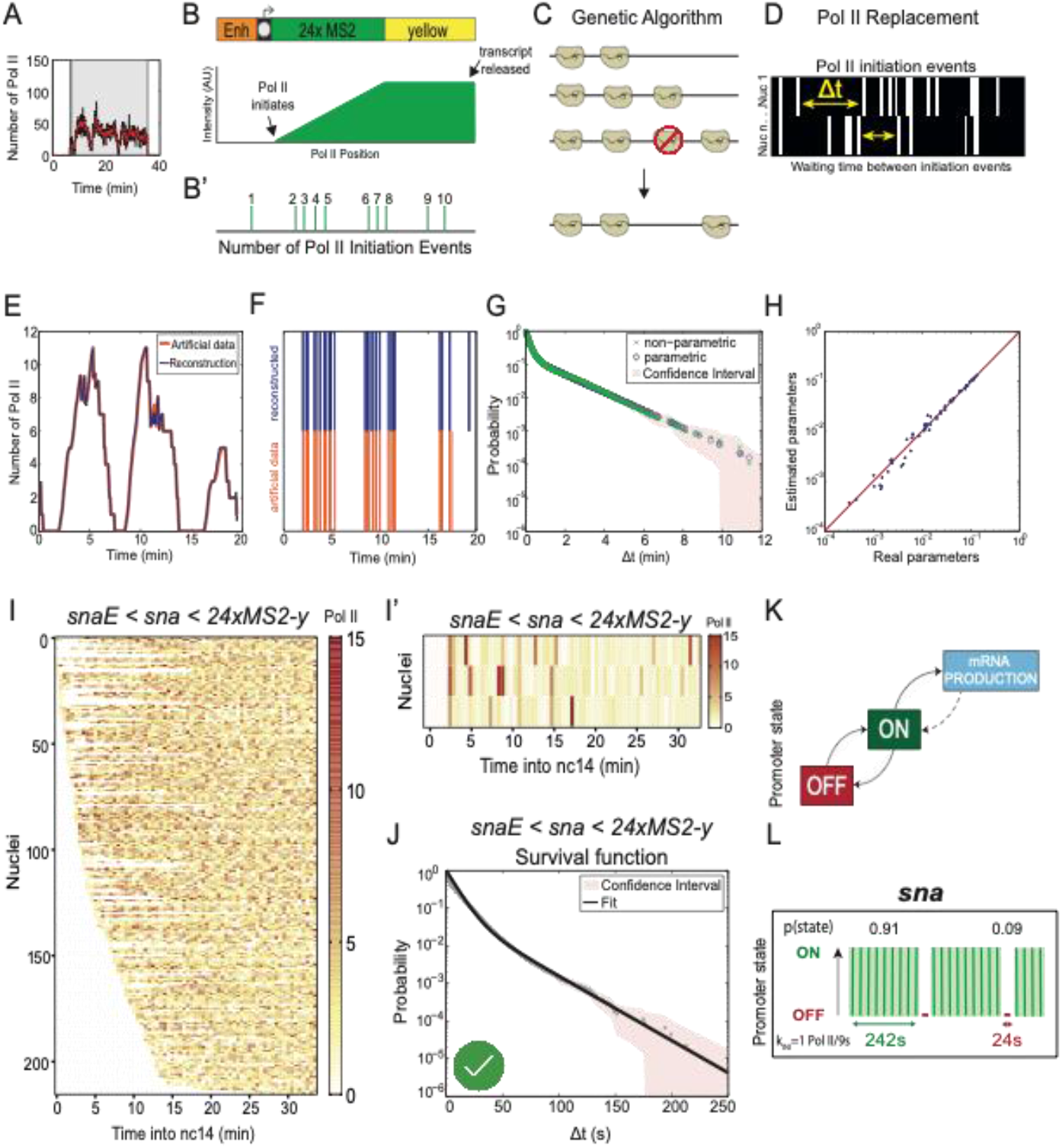
From live imaging of transcription to positions of initiation events, a machine-learning procedure. **A)** Representative trace of a single nucleus transcriptional activity. Grey box indicates the analysed transcriptional window, black curve represents the signal expressed as number of polymerases and red curve the reconstructed signal after the deconvolution procedure. **B)** Fluorescence intensity at the transcription site is a function of the passage of a single polymerase (B) as well as the number of polymerase initiation events (B’). **C)** A genetic algorithm is used to decompose fluorescence intensities and optimally locate polymerases within the gene body. **D)** For each nucleus, the position of polymerases is used to extract Δt, the lag time between two successive initiation events. **E)** Simulated representative trace (orange) and number of polymerases after deconvolution (blue). **F)** Known positions of polymerases (orange) and reconstructed polymerase positioning after deconvolution. **G)** Non-parametric (green) and parametric (blue) survival function estimates for simulated data. Dashed lines indicate 95% confidence interval. **H)** Comparison of known parameters used to generate artificial data (real parameter) with parameters obtained by applying our analysis pipeline (estimated parameters). Red line indicates perfect concordance. **I)** Heatmap showing number of polymerases for the 216 *snaE<sna* nuclei as a function of time. Each row represents one nucleus, and the number of Pol II initiation events per 30s bin is indicated by the bin colour. (I’) A representative population. **J)** Survival function of the distribution of waiting times between polymerase initiation events (red circles) and the two-exponential fitting of the population (black line, p-value KS test = 0.717). Red dashes represent 95% confidence interval (see **Supplemental Table 1**). Green check indicates accepted fitting. **K)** A two state model showing the probabilities to be in either the permissive ON state or the inactive OFF state for the *snaE<sna* transgene. **L)** Representation of estimated bursting dynamics for the *snaE<sna* transgene. Permissive ON state durations are depicted in green and inactive OFF states in red, and probabilities of each state shown above (see also **Supplemental Table 2**). **Statistics:** *snaE<sna*, 216 nuclei, 3 movies; see **Supplemental Movie S1**.

We then used a recently described novel machine-learning method to infer the transcriptional bursting mechanism^59^. This method involves three major steps: detection of successive initiation events for each nucleus, multi-exponential parametric regression of the distribution of waiting times between successive events, and identification of Markovian promoter state transition models.

In order to detect initiation events, we considered that each trace results from the convolution between i) the sequence of initiation events marked by a rise in GFP intensity plotted over time, and ii) the signal produced by a single polymerase (**Figure 2B, B’**)^33,59^. The deconvolution procedure uses a genetic algorithm to determine optimal Pol II positioning within the gene body (**Figure 2C**) and thus the time between successive Pol II initiation events (Δt) (**Figure 2D**). In this first step, some specific parameters are fixed, such as the temporal resolution (3.86s), the speed of Pol II elongation (45bp/s in *Drosophila* embryos^60^), the size of the transcript (5.6kb) and the retention time of the mRNA at the transcription site (assumed to be small relative to the time required to produce a transcript).

Having obtained temporal maps of individual Pol II transcription events, the method then statistically analyzes the distribution of waiting times between two successive polymerases (Δt)^59^. A multi-exponential regression fitting was applied to the distribution of Δt, indicating the number of rate-limiting transitions required to best fit the data (**Methods**). We used confidence intervals and a Kolmogorov-Smirnov test to rigorously determine the smallest number of rate-limiting steps fitting our experimental data (**Supplemental Table 1**). Following the principle of parsimony and to avoid overfitting, models with a larger number of steps and more parameters were not retained even if they also fit well. Importantly, this step uncovered the number of characteristic timescales of transcriptional fluctuations that is tantamount to the number of promoter states.

Finally, a transcriptional model of the promoter was established, with multiple states and timescales inferred from the parameters of the multi-exponential Δt distribution (**Supplemental Table 2**). This step permitted the estimation of the kinetic parameters such as the k_ON_ (rate of switching from a transcriptionally non-permissive state to a permissive one), k_OFF_ (switching rate from a transcriptionally permissive to a non-permissive state), and k_INI_ (the rate of Pol II initiation events once the promoter is in a transcriptionally permissive state) for the simplest two state model. However, kinetic estimates of more complex models (three or more promoter states) can also be derived from the Δt distribution. The accuracy and robustness of the deconvolution method was tested by applying it to artificial data with known positions of transcription initiation events and known kinetic parameters (**Figure 2E-H**). These artificial data were generated using the Gillespie algorithm and proved the robustness of the approach (see Methods).

Importantly, this approach provides a time map of transcription events in a cell population in a model-independent manner. The number of promoter states is evaluated during the multi-exponential fit procedure, but with no *a priori* constraints on the result^59^. This contrasts with current methods which directly fit a particular transcription model to the data, such as methods based on the autocorrelation functions^61–63^, maximal likelihood estimates^36^, or Bayesian inference^30,64,65^.

### The sna promoter as a model to investigate the impact of core promoter on transcriptional dynamics

To investigate the quantitative dynamics of the *sna* promoter, we applied this method to hundreds of nuclei pooled from *snaE*<*sna* embryos. A heatmap of positions of single initiation events for each nucleus revealed the *sna* promoter drove frequent transcriptional initiation events. Silent periods with no initiating polymerase were very rare (**Figure 2I-I’**, white bars), consistent with the results obtained from fixed embryos^21,66^. We found that a bi-exponential (sum of two exponential functions) fitting correctly fit the survival function of polymerase waiting times (**Figure 2J**). The bi-exponential fitting is only compatible with a two-state model (**Figure 2K**). Thus, transcriptional activity of the *sna* promoter can be described by the random telegraph model, with a simple random switch between an inactive OFF and a permissive ON state, from which transcription initiates at a given rate (k_INI_; **Figure 2K**). The probability to occupy the permissive ON state was high in the case of *sna* (0.91; **Figure 2L**). The T_ON_ was estimated to be 242s, with a T_OFF_ of 24s. The k_INI_ of *sna* promoter was estimated at one initiation event every 9s (**Figure 2L**), consistent with the inferred initiation rate at its endogenous locus^66^ and in the range of recent estimates of initiation frequencies for other developmental genes in *Drosophila*^32^ and in mammalian cells^37^. Collectively this shows that we can locate individual initiation events, determine the number of promoter states, select a promoter model by principle of parsimony, and then estimate average promoter switching rates *in vivo*.

### The TATA box regulates the ON and OFF duration but not the number of states

To determine how the TATA box influences transcriptional kinetics *in vivo*, we developed a series of *sna* core promoter mutants (**Figure 3A**) where the TATA box was replaced with either the TATA-like sequence of *kr* (*snaTATAlight*) that is bound by TBP^50^ (**Supplemental Figure 1**), or a non-TATA sequence where the first four bases are mutated (*snaTATAmut*). Surprisingly, the strong TATA mutation did not completely abolish the activity of the *sna* promoter, although this mutant promoter was devoid of any known core promoter motifs. In comparison to the *sna* promoter, the synchrony of the *sna* TATA mutants was reduced in a graded manner relative to the fidelity of the TATA box (**Figure 3B, Supplemental Movies S1-S3**). The *sna* promoter reached synchrony at approximately 11 minutes into nc14, while the *snaTATAlight* promoter reached synchrony at 24 minutes and the *snaTATAmut* promoter never reached synchrony. This implies that the TATA box influences synchrony of activation, most likely by promoting and stabilizing TBP binding (**Supplemental Figure 1**).

**Figure 3:**
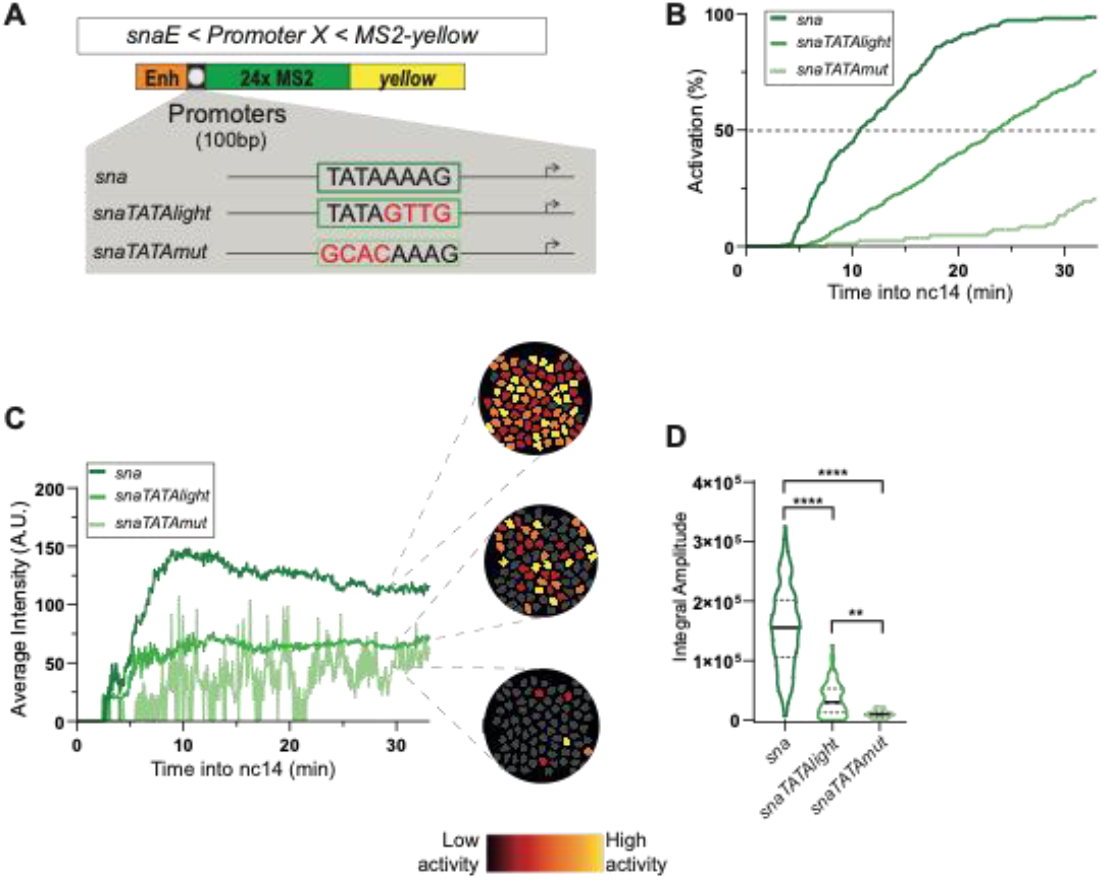
Decoding the role of the TATA box on promoter dynamics. **A)** TATA box mutations of the *sna* promoter. The *snaTATAlight* mutation corresponds to the sequence of the non-canonical TATA box from the *kr* promoter. **B)** Cumulative active nuclei percentage within mesodermal domain for indicated genotypes. **C)** Average instantaneous fluorescence of transcriptional foci of active nuclei during the first 30 minutes of nc14. Time zero is from anaphase during nc13-nc14 mitosis. False-coloured panels on right are coloured according to instantaneous fluorescence intensity, with inactive nuclei shown in grey. **D)** Distribution of individual trace integral amplitudes from first 30 minutes of nc14. Solid line represents median and dashed lines the first and third quartiles, using a Kruskal-Wallis test for significance with multiple comparison adjustment. ** p <0.01, **** p < 0.0001 **Statistics:** *snaE<sna*, 216 nuclei, 3 movies; *snaE<snaTATAlight*, 353 nuclei, 6 movies; *snaE<snaTATAmut*, 21 nuclei, 3 movies. See **Supplemental Movies S1-S3**.

Similar to the activation profiles, instantaneous TS intensities were directly correlated with TATA box sequence fidelity (**Figure 3C**). Interestingly, while both the *sna* promoter and *snaTATAlight* promoters appeared to reach a steady state, the *snaTATAmut* was unable to do so. This may have been the result of MCP-GFP signal being too low to consistently surpass the detection threshold, as single molecule FISH experiments indicated most nuclei in the ventral region were at least weakly active in nc14 *snaTATAmut* embryos (**Supplemental Figure 4A-C**). The total mRNA production of each promoter measured by the integral amplitude was similarly affected (**Figure 3D**), with the *sna* promoter driving highest total mRNA expression while the TATA box mutants expressed reduced amounts of mRNA correlated to the fidelity of the TATA box.

We next asked whether the *snaTATAlight* and *snaTATAmut* transgenes still followed a two-state model. For each nucleus of each of these three transgenic genotypes, we located single Pol II initiation events during the first 30 minutes of nc14 (**Figure 4A-F** and **Supplemental Figure 4D-F**). By examining the survival curve of Pol II waiting times, the dynamics of the *sna* TATA mutants appeared to be accurately recapitulated by the two-state random telegraph model, similarly to the unmutated *sna* promoter (**Supplemental Figure 4G-I, Supplemental Table 1**). We then analyzed which kinetic parameters changed in the mutant promoters. Extracting kinetic parameters for the *snaTATAlight* transgene revealed a relatively mild effect on initiation rates (1 Pol II/13s for *snaTATAlight* instead of 1 Pol II/9s) (**Figure 4G-I**, **Supplemental Table 2**). In contrast, we observed a strong increase in the T_OFF_ (**Figure 4G-K**) for the *snaTATAlight* promoter relative to the *sna* promoter (**Figure 4G-I,K, Supplemental Figure 5B**). The strong TATA mutation led to a similar trend, although due to the weak activity of this promoter, the number of analyzed nuclei for this mutant are lower than for the other promoters. The kinetic parameters of TATA mutant promoters led to an overall reduction in burst size, defined as the number of transcripts produced during an active period (**Figure 4L**). The reduction was primarily due to a decrease in the duration of the ON periods, consistent with a destabilized TBP/TATA box interaction.

**Figure 4:**
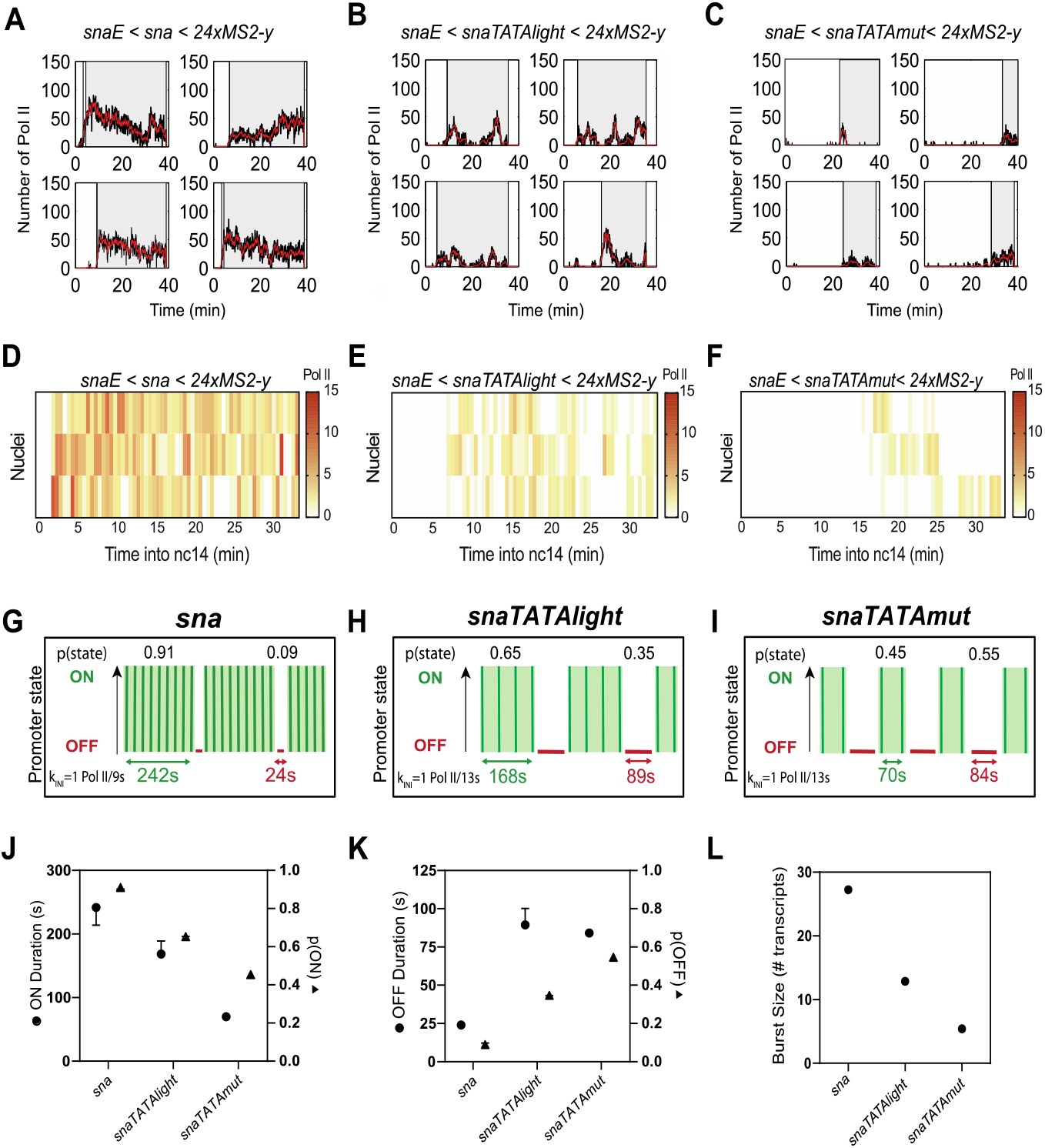
TATA box motif leads to long durations of a permissive promoter state. **A-C)** Representative traces of single nuclei transcriptional activities for the indicated genotypes. Grey boxes indicate the analysed transcriptional windows, black curves represent the signal expressed as number of polymerases and red curve the reconstructed signal after the deconvolution procedure. **D-F)** Heatmaps indicating number of polymerases during nc14 for each genotype. **G-I)** Representation of estimated bursting parameters for *sna* (G), *snaTATAlight* (H) and *snaTATAmut* (I) promoters (see also **Supplemental Table 1** and **Supplemental Table 2)**. **J)** Duration and probability of the permissive ON state. Error bars show the minimum/maximum of the error interval. **K)** Duration and probability of the OFF state. Error bars show the minimum/maximum of the error interval. **L)** Estimated burst size calculated as k_INI_/k_OFF_^75^ using the optimal kinetic parameters (**Supplemental Table 2**). **Statistics:** *snaE<sna*, 216 nuclei, 3 movies; *snaE<snaTATAlight*, 353 nuclei, 6 movies; *snaE<snaTATAmut*, 21 nuclei, 3 movies. See **Supplemental Movies S1-S3 and Supplemental Table 2**.

The impact of TATA on the duration of ON/OFF promoter states observed in *Drosophila* embryos is also in agreement with results from live imaging of *HIV-1* transcription in human cells^37^, and with genomic studies in cultured mammalian cells, where TATA-driven promoters are associated with large burst sizes^67^. They differ from those obtained with mutations of the *actin* promoter in *Dictyostelium*^36^ and this may be possibly due to differential behaviour between induced genes (developmental, HIV) and constitutive genes^36^.

We conclude that in *Drosophila* embryos, the TATA box largely controls gene expression through lengthening the ON state duration at the expense of the OFF state duration, but does not alter the number of promoter states.

### The INR motif induces a third rate-limiting promoter state

At the biochemical level, transcriptional activation is a multistep process requiring the orchestration of many factors^3^, and yet TATA-driven promoters can be modeled with a simple two state model corresponding to one kinetic rate-limiting step in this complex process. We hypothesized that additional rate-limiting steps were likely to exist and could be identified using our imaging-based methodology. As the TATA box did not affect the number of promoter states for the *sna* promoter, we examined the effect of another core promoter motif, the INR. In *Drosophila* cells and embryos, the INR is associated with stably paused genes, genome-wide ^7,9,15,68^. Moreover, analysis in cell culture indicated that genes with increased pausing stability tend to harbour an INR motif in the core promoter, and in particular a preferential G at the +2 position^12^. We therefore reasoned that manipulating the INR motif in embryos might affect pausing and therefore may induce changes in rate-limiting promoter states.

To examine the role of the INR core promoter motif, we created a series of transgenes in which we manipulated the INR without changing the enhancer or the downstream gene sequence (**Figure 5A**). The *sna* core promoter does not have an INR, while the core promoter of *kr* has a natural INR sequence with a G at +2 and is stably paused^16,18^. Moreover, the *kr* promoter possesses a non-canonical TATA box (*TATAlight*, **Figure 1A**); loss of both motifs leads to a non-functional promoter (data not shown). We exchanged the TSS region of *sna* with the INR of the *kr* promoter (*sna+INR*). We also created a transgene (**Figure 5A**) whereby the INR of *kr* was replaced with the TSS region of *sna* (*kr-INR1*). Interestingly, neither loss of the INR in the *kr-INR1* transgene nor gain in the *sna+INR* transgene strongly affected synchrony (**Figure 5B**, **Supplemental Movies S1, S4-S5**). Addition of an INR to *sna* did not dramatically affect mRNA output (**Figure 5D**). Both the *sna+INR* and *kr-INR1* transgenes had a higher instantaneous activity than the cognate wild type promoters (**Figure 5C**). Moreover, a second mutation of the INR in *kr* (*kr-INR2*) had similar synchrony and mRNA production (average instantaneous intensity) as in the wild type *kr* promoter (**Supplemental Figure 6A-C**). We then applied the deconvolution procedure to each genotype (**Figure 6A**, **Supplemental Figure 6D-K, Supplemental Figure 7A,D-K**). Surprisingly, our analysis of polymerase initiation events indicated that, for *sna+INR* nuclei, a two-state model was not sufficient to fit the data (**Figure 6A-B, Supplemental Figure 7I, Supplemental Table 1**). Instead, the *sna+INR* survival function was well fitted by adding an extra exponential term. Thus, a three-state model appropriately recapitulated *sna+INR* promoter dynamics (**Figure 6A-B, Supplemental Figure 7I-I’**). Similarly, the *kr* promoter also required a three-exponential fitting (**Figure 6B, Supplemental Figure 6H-H’**). In contrast, removing the INR from this promoter (*kr-INR1* promoter) led to a two-exponential fitting for the survival function (**Figure 6B, Supplemental Figure 6I-I’**). Similar results were obtained with a second INR mutant (*kr-INR2*) (**Supplemental Figure 6J-J’**). Furthermore, the natural INR-driven *Ilp4* promoter was also associated with a three-exponential fitting, and mutation of its INR sequence (*Ilp4-INR*) resulted in a reversion to a two-exponential fit (**Figure 6B, Supplemental Figure 8D-E,H-I**). These results therefore suggest that the INR motif is associated with a third rate-limiting promoter state.

**Figure 5:**
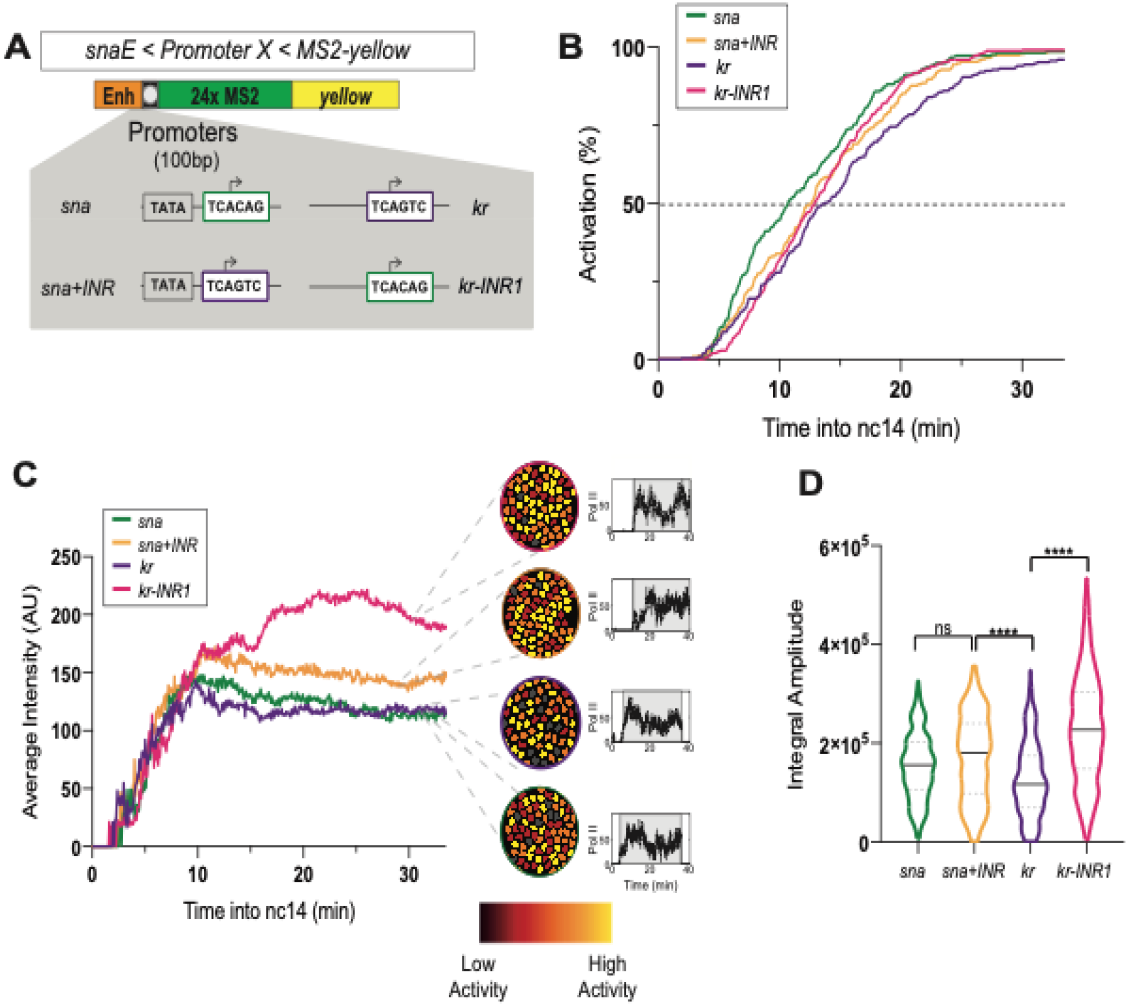
Decoding the role of the Initiator motif on promoter dynamics. **A)** Schematic of the transgenes used to decipher the impact of the INR motif. The *sna* core promoter has the TSS region replaced with the INR of *kr* (*sna+INR*), and the *kr* core promoter INR motif is replaced with the TSS region of *sna* (*kr-INR1*). **B)** Cumulative active nuclei percentage within mesodermal domain for indicated genotypes. **C)** Average instantaneous fluorescence of transcriptional foci of active nuclei during the first 30 minutes of nc14. Time zero is from anaphase during nc13-nc14 mitosis. False-coloured panels on right are coloured according to instantaneous fluorescence intensity with inactive nuclei in grey. **D)** Distribution of individual trace integral amplitudes from first 30 minutes of nc14. Solid line represents median and dashed lines the first and third quartiles, using a Kruskal-Wallis test for significance with multiple comparison adjustment. ** p <0.01, **** p < 0.0001 **Statistics:** *snaE<sna*, 216 nuclei, 3 movies; *snaE<sna+INR*, 236 nuclei, 4 movies; *snaE<kr*, 243 nuclei, 4 movies; *snaE<kr-INR1*, 342 nuclei, 5 movies. See **Supplemental Movies S1, S4-S6**.

**Figure 6:**
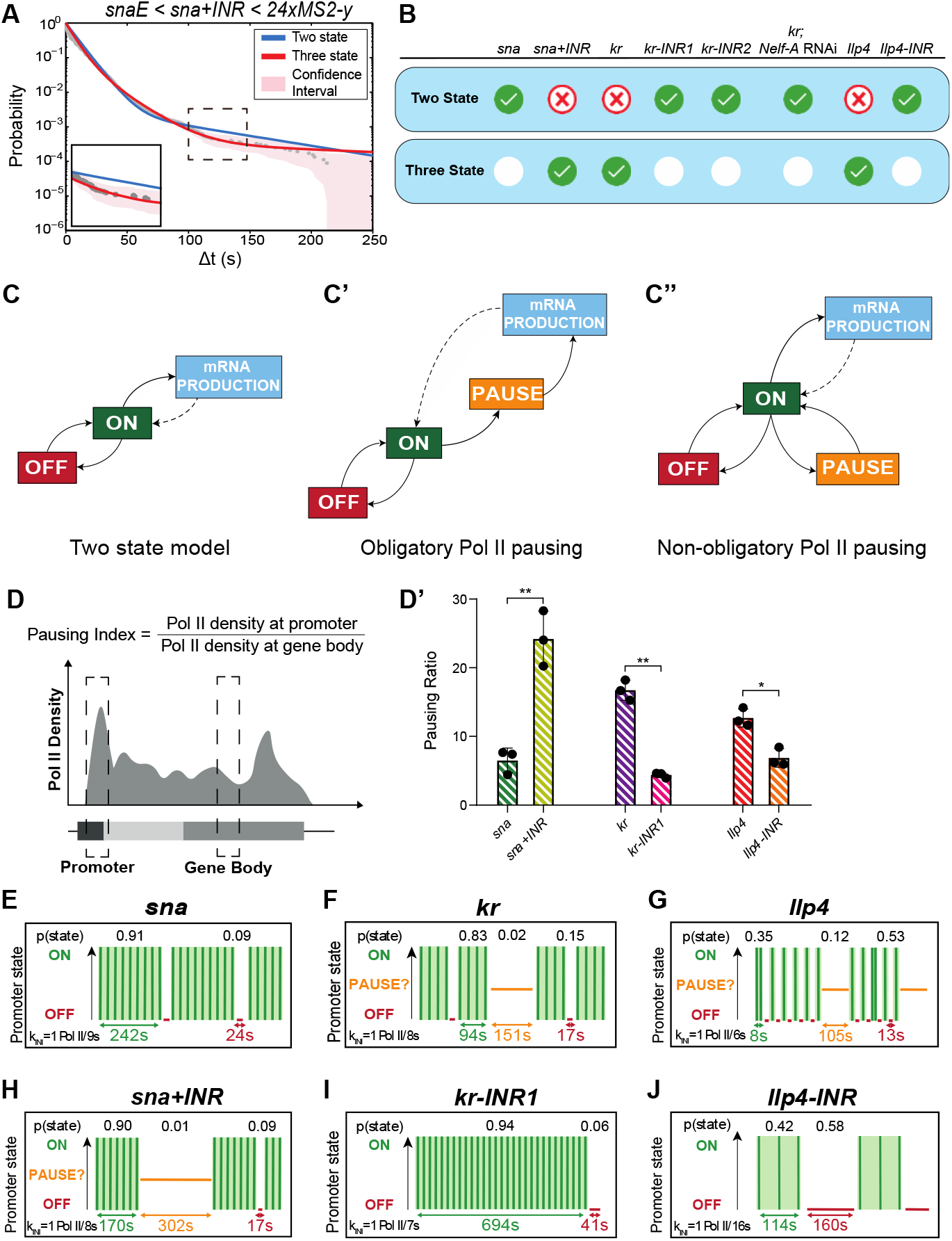
The INR motif induces an extra promoter state related to pausing. **A)** Survival function of the distribution of waiting times between polymerase initiation events (grey circles). Two-exponential fitting of the population (blue line, p-value KS test = 1.37e^−4^), and three exponential fitting (red line, p-value KS test = 0.999). Shaded region represent 95% confidence interval (see also **Supplemental Table 1)**. **B)** Table indicating the most parsimonious number of states required to fit each indicated genotype (see also **Supplemental Table 1)**. Green check indicates an accepted fitting. **C)** Schema of the two-state telegraph model (C) a three-state model with obligatory pausing (C’) and a three state model with non-obligatory pausing (C’’). **D)** Pausing index calculation (D) and pausing index of indicated transgenes (D’) measured by ChIP-qPCR (average + SD). n = 3 biological replicates. ** p < 0.01, * p < 0.05 with a Mann Whitney test. **E-J)** Representation of *sna*, *sna+INR*, *kr*, *kr-inr1, Ilp4 and Ilp4-INR* estimated bursting dynamics. Permissive ON states are in green, inactive PAUSE states in orange, and inactive OFF states in red. **Statistics:** *snaE<sna*, 216 nuclei, 3 movies; *snaE<sna+INR*, 236 nuclei, 4 movies; *snaE<kr*, 243 nuclei, 4 movies; *snaE<kr-INR1*, 342 nuclei, 5 movies, *snaE<snaTATAlight*, 353 nuclei, 6 movies; *snaE<snaTATAlight+INR*, 193 nuclei, 3 movies; *snaE<Ilp4,* 114 nuclei, 2 movies*; snaE<Ilp4-INR*, 48 nuclei, 2 movies. See **Supplemental Movies S1, S4-S6, Supplemental Table 2**.

### Pol II pausing is associated with a third promoter state

Given the correlation between the presence of an INR motif and the number of characteristic timescales and thus number of promoter states (three in the presence of an INR), we reasoned that one of these states could be polymerase pausing. To test this hypothesis, we impaired the establishment of pausing by reducing the expression of the largest subunit of the pause-inducing Negative Elongation Factor complex (NELF), NELF-A. Previous work demonstrated that depletion of NELF by RNAi globally reduced promoter proximal pausing in *Drosophila* cells^13^, and that loss of the human C-terminal NELF-A “tentacle” blocked stabilization of pausing^69^. We combined the inducible RNAi/GAL4 system with our MS2/MCP-GFP reporter to examine transcriptional dynamics of the three state *kr* promoter. We performed live imaging of the *kr*-MS2 transgene and compared a control *white* RNAi knockdown with embryos harbouring a maternal depletion of *Nelf-A* (evaluated to >85% knock-down by RT-qPCR, **Supplemental Figure 9A**). Remarkably, reducing *Nelf-*A transcript levels led to a change in *kr* promoter dynamics, with a reversion to a two-promoter state dynamic and much less frequent long waiting times (**Figure 6B** and **Supplemental Figure 9E-F**, arrowheads, **Supplemental Table 1**). These data, combined with the effect of INR mutations in *cis*, led us to propose that one of the two inactive promoter states observed with the three state promoters could be associated with promoter-proximal polymerase pausing.

To confirm that the INR motif was truly altering Pol II pausing, we performed Pol II ChIP-qPCR on our promoter transgenes from staged embryos. We evaluated the strength of pausing by computing the pausing index. The pausing index (PI) was obtained by quantifying the Pol II binding at the promoter relative to the gene body^70^ (**Figure 6D**). As shown previously, in our transgenic embryonic extracts encompassing various stages from nc13 to late nc14, the wild type *sna* promoter is paused ^18,21,50^, but the addition of an INR motif significantly increased its PI (**Figure 6D’**), consistent with INR-mediated lengthening of the pause duration observed in cell culture^12^. To examine the inverse scenario, we also quantified the PI of the *kr* and *kr-INR1* transgenic promoters. Similar to what is observed in the endogenous promoter^18^, the *kr* transgene was highly paused (**Figure 6D’**). Mutation of the INR in the *kr-INR1* transgene reduced pausing (**Figure 6D’**). Likewise, the *Ilp4* promoter from our transgene showed a high level of pausing, which decreased upon mutation of its INR sequence (**Figure 6D’**). Both the *sna+INR* and *kr* promoters also show increased NELF-E enrichment at these promoters by ChIP-qPCR on embryonic extracts (**Supplemental Figure 9G**). Taken together, characterization of the INR core promoter motif suggests the presence of a strong INR motif results in stabilization of pausing *in vivo*.

### Pol II does not undergo systematic pausing

Next, we envisaged how pausing could be modeled by three distinct promoter states. The current view of pausing is that all polymerases systematically enter into pausing, a step followed by either a productive elongation or a termination^17,71,72^. We therefore initially tested if our data could be explained by a model consisting of three promoter states (inactive OFF, permissive ON, and paused) where all productive polymerases undergo pausing prior to transitioning to the permissive ON state (**Figure 6C’**). This model was clearly incompatible with our data, as the fitting exceeded the bounds of the 95% confidence interval (**Supplemental Figure 6L, Supplemental Figure 7I”**). Recent work on the prototypic model of a highly paused promoter, the *HIV-1* promoter, also found that modeling transcriptional dynamics with obligatory pausing was not in agreement with live imaging data in HeLa cells, and instead proposed an alternative model of pausing where only a fraction of polymerases are subject to pausing in a stochastic manner^59^. We asked if this alternative non-obligatory pausing model could be applicable to our findings (**Figure 6C’’**). According to the goodness of fit, non-systematic pausing was compatible (**Supplemental Figure 6H’** and **Supplemental Figure 7I’**). Our machine-learning method enabled the estimation of the kinetic transition rates between these states, summarized in **Supplemental Table 2** and **Supplemental Figure 5A-C**.

We conclude the presence of an INR motif translates into a longer paused state that creates an additional rate-limiting step during transcription *in vivo*. In our kinetic model, the inactive states could in principle equally correspond to pausing. These two promoter states are discernible by their distinct timescales, on the order of seconds and minutes respectively (**Supplemental Table 2**). Because the NELF knock-down dramatically decreases the frequency of long waiting times (**Supplemental Figure 9E-F**), we favour the longer-lived inactive state, which lasts on the order of minutes, as the potential pausing state. The second short-lived inactive state, lasting on the order of seconds and present for both TATA- and INR-containing promoters, would then correspond to a non-permissive promoter state, possibly TFIID-unbound as its frequency increases in TATA mutants (**Figure 4K, Supplemental Table 2**). As the *sna* and *kr* endogenous promoters drive a high level of gene expression (**Supplemental Figure 1**), it is reasonable to hypothesize that such a non-permissive state is transient and infrequent (**Supplemental Table 2**).

### INR control of promoter switching kinetics

Like the natural *sna* promoter, the *sna+INR* promoter showed a high probability of occupying the transcriptionally permissive ON state, with a long lifetime of 170s. This ON duration was slightly reduced compared to the *sna* transgene (**Figure 6E,H**), possibly implying competition between the natural canonical TATA box and the added INR in the *sna+INR* combination (see below). The major difference between these two promoters was the existence of two distinguishable inactive states linked to the presence of the INR sequence. One of these was short-lived (17s) and one long-lived (302s) (**Figure 6E-H**, **Supplemental Figure 5B-C, Supplemental Table 2**). The stable long-lived paused state was not observed with the *sna* promoter but did occur in the presence of the INR-containing *kr* promoter.

In the case of *kr* and *Ilp4* developmental promoters, the paused state lasted approximately 151s and 105s respectively but was reached relatively rarely (**Figure 6F-G**, **Supplemental Figure 5C, Supplemental Table 2**). Pausing was not observed upon two types of mutations of the *kr* INR motif (**Figure 6I, Supplemental Figure 6I-J,M, Supplemental Table 2**). Thus our data favours a model where the paused state lasts several minutes (in the case of *kr*, *sna+INR* and *Ilp4*), but occurs infrequently. Interestingly, the s*na* promoter is also moderately paused in our ChIP assay and at its endogenous locus^74^, but this promoter is not regulated by a three promoter state dynamic. We hypothesize that pausing is highly unstable for this promoter at this stage and does not constitute a rate-limiting step that we can capture with our live-imaging assay (see discussion).

In addition to two separable inactive states, the *kr* transgene exhibited a highly probable permissive state (0.83) with a duration of 94s (**Figure 6F, Supplemental Figure 5A, Supplemental Table 2**). When the INR of *kr* was mutated (*kr-INR1*), there was a significant increase in ON duration (**Figure 6I**, **Supplemental Table 2**). A similar effect was obtained with an alternative mutation of the INR (*kr-INR2*) (**Supplemental Figure 5A, Supplemental Figure 6M, Supplemental Table 2**).

Interestingly, in all the transgenes, the differences in the number of states and the active/inactive durations were not accompanied by a change in Pol II firing rates. The *sna*, *sna+INR*, *kr* and *kr-INR1* mutant all exhibited an estimated k_INI_ of 1 Pol II every 8-9 seconds (**Figure 6E-F,H-I, Supplemental Table 2**). These estimates agree with recently documented polymerase initiation rates of *Drosophila* developmental promoters^30,32,75^. Our results suggest the rate of initiation is not associated with the presence or absence of an INR motif. Instead, the INR motif leads to a regulation of transcriptional bursting via two inactive promoter states, one of which is associated with stabilized pausing.

Taken collectively, our results demonstrate that transcription dynamics are differentially regulated by the INR and the TATA box. Developmental promoters containing a strong INR motif can travel through multiple inactive states due to an extra regulatory step at early elongation via Pol II pausing. Our live imaging data favour a model whereby pausing occurs on the order of minutes but is not an obligatory state reached by all engaged polymerases. Instead, we propose that when pausing constitute a rate-limiting step, it occurs for only a subset of polymerases with a low frequency.

### Promoter sequences and timing of initiation

A second property that we found to be common to all promoters examined, was the regime of waiting times between mitosis and first transcriptional activation. The analysis of the lag time between mitosis and onset of activation is ‘non-stationary’ and was previously modeled using a different modeling paradigm, based on mixed gamma distributions of the random time to transcription activation after mitosis in nc14^57^. This analysis estimates two parameters, the number of promoter states (a) and transition times between them (b). We applied this modeling framework to our data by only focusing on the distribution of waiting times between mitosis and first activation (illustrated in **Supplementary Figure 10A**). Interestingly, a multi-exponential distribution becomes a mixed gamma distribution when the time parameters of the exponentials are even. Conversely, the equality of these time parameters justifies the use of a mixed gamma regression. Using the more general multi-exponential regression, we proved that regardless of the promoter sequence, the transition times between states during this lag regime are homogeneously distributed (**Supplementary Figure 10B-C**). This is in contrast with the transition times between states in the stationary bursting regime which are heterogeneous. This suggests that the states and transitions involved in the two regimes (non-stationary and stationary) are distinct and have separate regulation. This further supports a previous study that demonstrated the delay in post-mitotic transcription activation is dependent on the enhancer sequence^57^.

### Interplay between TATA and INR

Recent single cell RNA-seq quantification suggests that promoters exhibiting both TATA and INR motifs produce higher burst sizes than when only one motif is present^67^. However, a systematic interrogation of human promoters with a synthetic biology approach shows that TATA and INR additively but not synergistically increase gene expression^76^. A synergistic and/or additive effect does not seem to be evident in *Drosophila*, as TATA-containing promoters have relatively infrequent INR motifs or exhibit and INR sequence devoid of a G at +2, shown to be associated with long pause durations^12,50^.

To quantify TATA box and INR interplay on rate-limiting promoter switching states *in vivo*, we compared the *sna+INR* and *snaTATAlight* transgenes to that of a combined mutant *snaTATAlight+INR* promoter (**Supplemental Figure 7A**). Remarkably, after applying the deconvolution procedure to *snaTATAlight+INR* nuclei (**Supplemental Figure 7G**), the distribution of waiting times between initiation events required a fit with two exponentials similarly to *snaTATAlight* (**Supplemental Figure 7K-K’**). We note however that the statistical tests allocating the number of promoter states for this particular genotype were not robust (**Supplemental Table 1**). In terms of promoter kinetics, the *snaTATAlight+INR* behaved similarly to the *snaTATAlight* promoter (**Supplemental Figure 5A-B**). Interestingly, the *snaTATAlight+INR* has a p(OFF) nearly twice that of the *snaTATAlight* promoter (**Supplemental Table 2**), indicating that the INR may act to maintain a non-permissive promoter state even in the absence of polymerase pausing.

To gain more insight into TATA/INR interplay, we also generated a *kr* mutant promoter that does not contain a non-canonical TATA (*kr-TATA*; **Supplemental Figure 6A**). Data from this promoter required a three-state model (**Supplemental Figure 6K-K’**), similar to the *kr* promoter. In the *kr-TATA* promoter, the duration and probability of the paused state increased while the ON state appeared shorter and less probable (**Supplemental Figure 5A,C** and **Supplemental Table 2**). This indicates that the loss of TBP binding may increase pausing induced by the INR motif, potentially providing a mechanistic rationale for the relative scarcity of dual TATA/canonical INR promoters^12,50^ in the *Drosophila* genome. Taken together, these results indicate that TATA and INR are not simply synergistic or additive but display more complex interactions.

## Discussion

The spatiotemporal organization of gene expression is critical to the development of a functional living organism. While we have accumulated knowledge on how enhancers precisely regulate gene expression, we know relatively little about the impact of core promoters on transcriptional bursting in a developing embryo. Here we investigated how minimal 100bp sequences of developmental promoters control transcriptional states and their switching kinetics. We found that a classical two state model does not suffice to capture promoter dynamics of stably paused promoters containing a strong INR motif but does fit with TATA-containing promoters.

### Promoter sequences and promoter states

The development of a novel numerical deconvolution method revealed startling insights into the variation of transcriptional initiation between minimal promoter sequences *in vivo*. Our experimental set-up allowed us to unmask gene expression variations, which are only dictated by variations in core promoter sequences. Indeed, cell cycle duration and synchrony, concentration of input transcription factors and chromatin states were intentionally kept constant. Our assay revealed that a similar transcriptional activity profile could be obtained from two very distinct promoter sequences via distinct modes of transcriptional initiation. While *sna* promoter dynamics could be explained by a simple two state model, the *kr* promoter required a three-state model, with two distinct inactive promoter states, a short lived and a longer lived one. These distinct transcriptional regimes are unlikely due to differences in enhancer-promoter specificity, as evidenced by the synchronous and high transcriptional activation reached with these two promoters. However, knowing the widespread preferential promoter code for specific enhancers^77,78^, it will be interesting to use our pipeline to examine which rate-limiting step of promoter dynamics is tuned by enhancer-promoter choice.

How could such a small number of states be compatible with the numerous promoter-binding events occurring during transcription initiation? Biochemical studies reveal structural states, while imaging-based approaches unmask key rate-limiting kinetic states. Whole genome methylation footprinting disentangled five transcription initiation states in *Drosophila* cultured cells^15^. Remarkably, the authors demonstrated that TATA-containing promoters were frequently found in PIC-bound configuration (PIC alone or PIC+Pol II). We therefore propose that the permissive state in *Drosophila* embryos corresponds to a state where promoter DNA is bound by the PIC. The inactive state, which we found to be very transient, could then possibly represent a TBP-unbound configuration consistent with TBP dynamics observed in human cells^37,39,79^.

### Promoter code, dynamics and zygotic genome activation

By using the *sna* promoter as a model, we found that TATA box directly impacts promoter occupancy of the active state and allows large transcriptional bursts by promoting long ON durations. How could the presence of a TATA box permit these long ON durations? Slow TBP protein turnover at human TATA box-containing genes may foster stable long ON durations^37,39,79^. However, this hypothesis needs to be confirmed by quantification of TBP kinetics during *Drosophila* ZGA.

In its endogenous context, the *snail* promoter is among the first genes to be transcribed during ZGA, in a particularly constrained environment with extremely short cell cycles (<15 minutes). Remarkably, the majority of genes expressed during this critical period are highly expressed, short, and intron-less with a canonical TATA box and are generally non-paused^18,50^. Subsets of these, including *snail,* are considered ‘dual promoters’, as they gradually acquire pausing as development proceeds^50^. Thus, it is possible that the kinetic bottlenecks regulating transcription of developmental promoters evolve as developmental timing proceeds, with paused polymerase gradually emerging as a rate-limiting step. In our study, the promoter model *sna* shows a moderate level of pausing (as measured by ChIP) but is regulated by a simple two-state model in nc14. This could indicate that pausing occurs on a timescale indistinguishable from the OFF state and is thus embedded within it. Strikingly, early transcription in zebrafish embryos occurs in similarly constrained rapid cell cycles where a subset of zygotically-activated genes also display a T/A-rich WW-box motif^80^. Thus, regulation of transcription via a unique rate-limiting step (OFF to ON transition) that is TBP-dependent might be conserved between fly and vertebrate embryos.

In this work, we provided quantitative evidence that INR containing promoters are associated with a three-state model (ON, paused, OFF), contrasting with TATA promoters.

Interestingly, genomics studies revealed that transcription initiation code evolves during ZGA in flies and vertebrates^50,54,80^. Given the results of this study and those obtained in *Drosophila* cultured cells^12,16^, it is tempting to link the gradual emergence and stabilization of pausing during ZGA with the presence of an INR, in particular an INR with a G at +2 position^12^. In light of our results, we propose that the switch in promoter usage from TATA-driven to INR-driven during ZGA may lead to a change in transcriptional dynamics from two to three states, to include an elongation-mediated checkpoint. Acquisition of an extra rate-limiting step might help control cell-to-cell expression variability as well as fine-tune gene expression levels.

Taken collectively, this study establishes our promoter imaging assay and novel mathematical deconvolution and modeling methodology as a valuable tool to probe gene expression dynamics during development. Quantitative analysis of promoter dynamics at high temporal resolution opens the door to a deeper insight into the molecular mechanisms underlying transcriptional regulation *in vivo*. Future studies involving direct manipulation of pause initiation or duration in living embryos using approaches such as optogenetics^81^ combined with this framework will help to establish a broader understanding of the nature of promoter states and the role of Pol II pausing *in vivo*.

## Supporting information

Supplemental Movie S1 - snaE<sna

Supplemental Movie S1 - snaE<snaTATAlight

Supplemental Movie S1 - snaE<snaTATAmut

Supplemental Movie S1 - snaE<sna+INR

Supplemental Movie S1 - snaE<kr

Supplemental Movie S1 - snaE<kr-INR1

Supplemental Movie S7 - nos-GAL4 < RNAi white, snaE < kr

Supplemental Movie S7 - nos-GAL4 < RNAi Nelf-A, snaE < kr

## Author contributions

Conception and design ML; Acquisition of data: MD, VP and CF; Analysis: ML, MD, VP and OR; Software AT, OR; Modeling: OR; Interpretation of data ML, MD, VP and OR; Writing: VP and ML with input from OR and EB; Visualization: MD, VP, OR; Supervision: ML; Fund acquisition: ML. OR and EB conceived and developed the machine learning deconvolution procedure and the facultative model of paused polymerase. All authors read the manuscript.

## Acknowledgements

We thank Julia Zeitlinger, Jean-Christophe Andrau, Jeremy Dufourt and all members of the Lagha lab for their critical reading of the manuscript and constructive discussions. We are grateful to F. Juge for insightful discussions and H. Faure-Gautron for technical assistance. We acknowledge the MRI imaging facility, a member of the national infrastructure France-BioImaging supported by the French National Research Agency (ANR-10-INBS-04, «Investments for the future»). This work was supported by the ERC SyncDev starting grant to M.L and CNRS; by CNRS grant PEPS MIGHTY to OR and EB, and by ANRS grants ECTZ62561 to EB. OR acknowledges support from the French National Research Agency (ANR-17-CE40-0036, project SYMBIONT). Machine learning calculations were performed on the HPC facility MESO@LR run by the University of Montpellier.

## Materials and Methods

### Drosophila stocks and genetics

All crosses were maintained at 25°C. Transgenic lines were maintained as homozygous stocks. For live imaging, homozygous males carrying the transgene of interest were crossed with homozygous females bearing the *MCP-eGFP-His2Av-RFP* construct. For smiFISH, homozygous flies were crossed to *yw* in order to facilitate single molecule detection. *nos*:GAL4-VP16 was recombined with UAS:*MCP-eGFP-His2Av-RFP* as previously described^57^. UAS:*white RNAi* (#35573) and UAS:*Nelf-A RNAi* (#32897) were obtained from the Bloomington Stock *Drosophila* Centre (University of Indiana, Bloomington IA).

### Cloning and Transgenesis

The *sna* distal enhancer-24xMS2-*y* mini-gene was previously described^51,57^. Promoters (**Supplemental Figure 1, Supplemental Table 3**) were amplified from genomic DNA using Q5 polymerase (New England Biolabs) and inserted between the enhancer and 24xMS2 sequences using restriction enzyme-mediated ligation. Mutations were performed with the QuikChange II Site Directed Mutagenesis kit (Agilent Technologies) or synthesized (Twist Biosciences) and inserted using restriction-mediated cloning. All constructs were sequenced to ensure appropriate insertion. Transgenic flies were generated using PhiC31-mediated recombination (Best Gene, Inc.), and all constructs were inserted into the same genetic background and genomic position (*BL 9750*).

### Live Imaging

Embryos were permitted to lay for 2h prior to collection for live imaging. Embryos were hand dechorionated and mounted on a hydrophobic membrane prior to immersion in oil to prevent desiccation, followed with addition of a coverslip.

Live imaging was performed with an LSM 880 with Airyscan module (Zeiss). Z-stacks comprised of 30 planes with a spacing of 0.5μm were acquired at a time resolution of 3.86s/stack in fast Airyscan mode with laser power measured using a ThorLabs PM100 optical power meter (ThorLabs Inc.), and maintained across embryos at at 3.8μW for constitutive MCP-GFP/His2A-RFP expression, and 5.0μW for RNAi analyses. All movies were performed with the following settings: GFP excitation by a 488nm laser and RFP excitation by a 561nm were captured on a GaAsP-PMT array with an Airyscan detector using a 40x Plan Apo oil lens (NA = 1.3) and a 3.0 zoom on the ventral region of the embryo centred on the presumptive ventral midline. Resolution was 512×512 pixels with bidirectional scanning. Airyscan processing was performed using 3D Zen Black (Zeiss).

### Live Imaging Analysis

Visualization and analysis of the time series were performed using a custom-made software developed in Python™ that permits visualization of each analysis step and manual correction if necessary (**Supplemental Figure 2**). Activation time traces were collected starting from Airyscan-processed Z-series described above. Green (MS2) and red (His2Av) channels were clipped to consider only after the start of nc14 as defined by the progression of anaphase across the region of interest. Nuclei were maximum intensity projected and pre-smoothed with a Gaussian filter and then thresholded with an Otsu threshold value. The resulting connected components of the binary images were then labeled and touching nuclei segmented with a watershed algorithm. The software enabled manual correction of the segmentation. Nuclei were finally tracked across the time frames using a minimum distance criterion plus a user defined distance threshold. Nuclei appearing for only a few frames or those touching the border were removed from the analysis to exclude sources of errors.

GFP puncta representing transcription sites were analyzed in 3D. Because the duplication of DNA occurs relatively early in nc14 and because of immediate proximity of sister chromatids ^30,82^, it is challenging to independently resolve individual sister chromatid signal using live imaging. This is why, the GFP signal at each transcriptional spot can be considered as the sum of both sister chromatids that we treat as a single transcriptional trace. For each time frame, the 3D image was filtered with a 3D Laplacian of Gaussian filter and then thresholded. The threshold value (THR) was expressed as μ + THR * σ, where μ and σ are the average and the standard deviation of the pixels values of the filtered image respectively, while THR is a user-defined value. Threshold value was in this way rescaled with respect to statistical properties of the filtered image, making THR a value independent of the particular data acquisition. All detected spots were filtered to remove 1) all the spots with a volume less than a user-defined volume threshold, and 2) spots present in only one z-frame. For each time frame, detected spots were associated in 2D to the overlapping or closest nucleus, inheriting the tracking from them; a user-defined distance threshold between nucleus and spot was used in order to avoid mis-associations.

Finally, each nuclei-puncta pair passing filtering was described as a time series of intensity, volume and position. To eliminate intensity variation within the Z-series, spot intensity values were divided by the background fluorescence of the average intensity value of the pixels surrounding the independent spots. Analysis was restricted to the region 25μm on either side of the centre of the gastrulation furrow present at the end of nc14 as positioned using a maximum intensity tile-scan of the entire embryo to determine coordinate position of the Z-stack. From this data, it is possible to extract: the timing of activation measured as the first timepoint GFP fluorescence crosses the software detection threshold for >1 Z-stack; cumulative activation; intensity profiles for individual nuclei; and individual nuclei burst trajectories. Integral amplitude was calculated using R to determine the surface area under the curve of each individual nuclei and analyzed with Prism (Graphpad) using a Kruskal-Wallis test for significance with multiple comparison adjustment. The calibration method has a minimum detection threshold of >3 transcripts per transcription site. All figures report the number of nuclei and movies used for analysis in figure legends. Movies of genotypes not supplied as supplemental movies available upon request.

### smiFISH

Embryos heterozygous for the transgene of interest were fixed in 10% formaldehyde/heptane for 25 minutes with shaking before a methanol quench and stored at −20°C in methanol before use. smiFISH probes targeting the *yellow* (*y*) reporter gene were designed as previously described ^83^ using FLAP-Y for secondary probe recognition (Integrated DNA Technologies, Inc.). Secondary probes were conjugated to Cy3 at the 5’ and 3’ ends (Integrated DNA Technologies, Inc.). Probes were resuspended in TE at appropriate equimolar concentration. Prior to probe addition to embryos, the *y* targeting primary probes were hybridized to the secondary FLAP-Y probes as described^83^ and maintained at −20°C in the dark prior to use.

Embryos were prepared for smiFISH as briefly follows: embryos were dehydrated with 2×5min washes in 100% ethanol, followed by rehydration in PBT for 4×15 minutes and equilibration in 15% formamide/1xSSC for 15 minutes. During equilibration, the smiFISH probe mixture was prepared with a final concentration of 1xSSC, 0.34 μg/μL *E. coli* tRNA (New England Biolabs), 15% formamide (Sigma), 5μL probe duplex, 0.2μg/μL RNAse-free BSA, 2mM vanadyl-ribonucleoside complex (New England Biolabs), and 10.6% dextran sulphate (Sigma) in RNAse-free water. The equilibration mixture was removed and replaced with probe mixture, and embryos were incubated overnight in the dark at 37°C. The following day, embryos were rinsed twice in equilibration mix and twice in PBT, followed with DAPI staining and three PBT washes before mounting in ProLong Gold mounting media (Life Technologies).

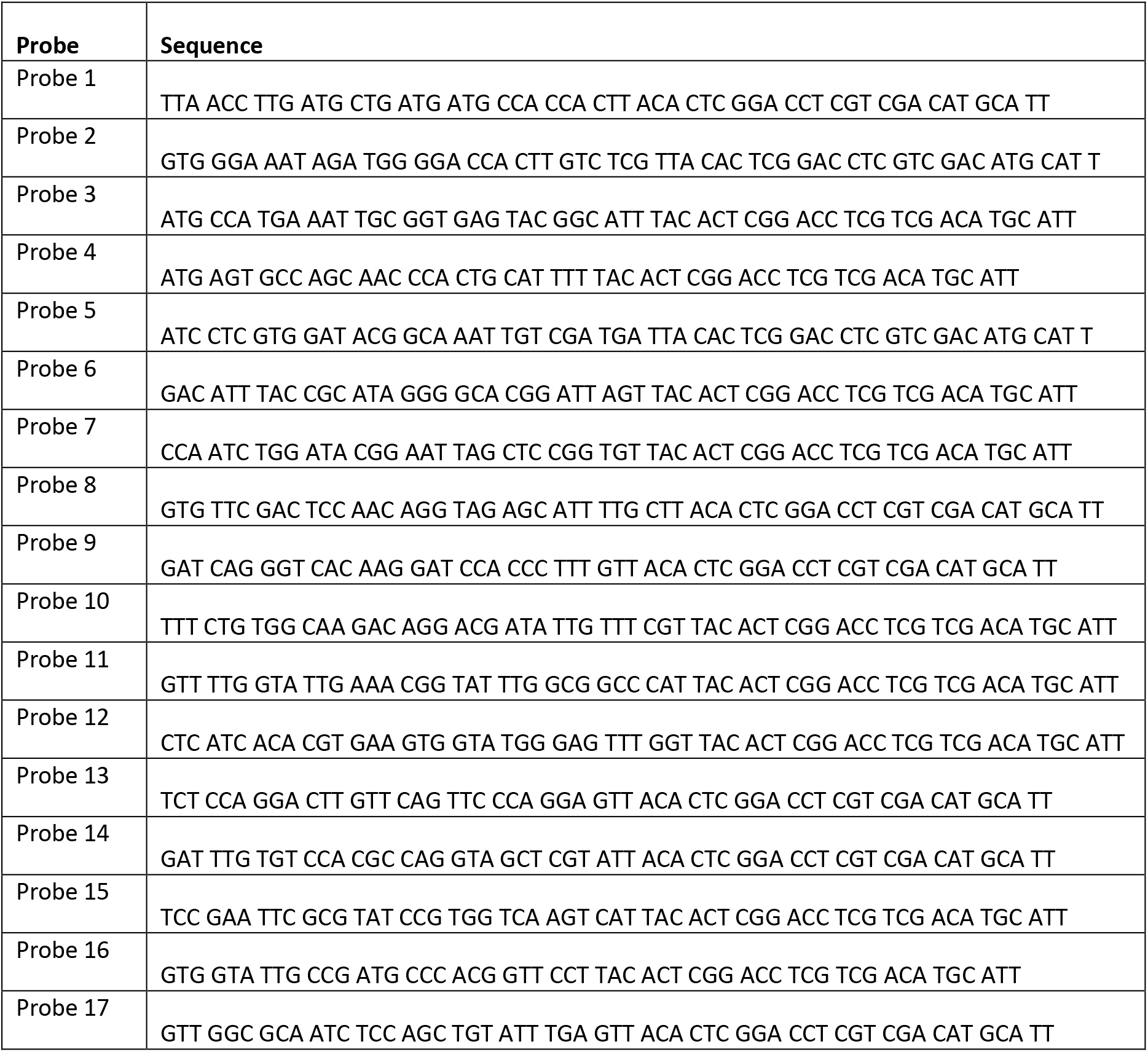

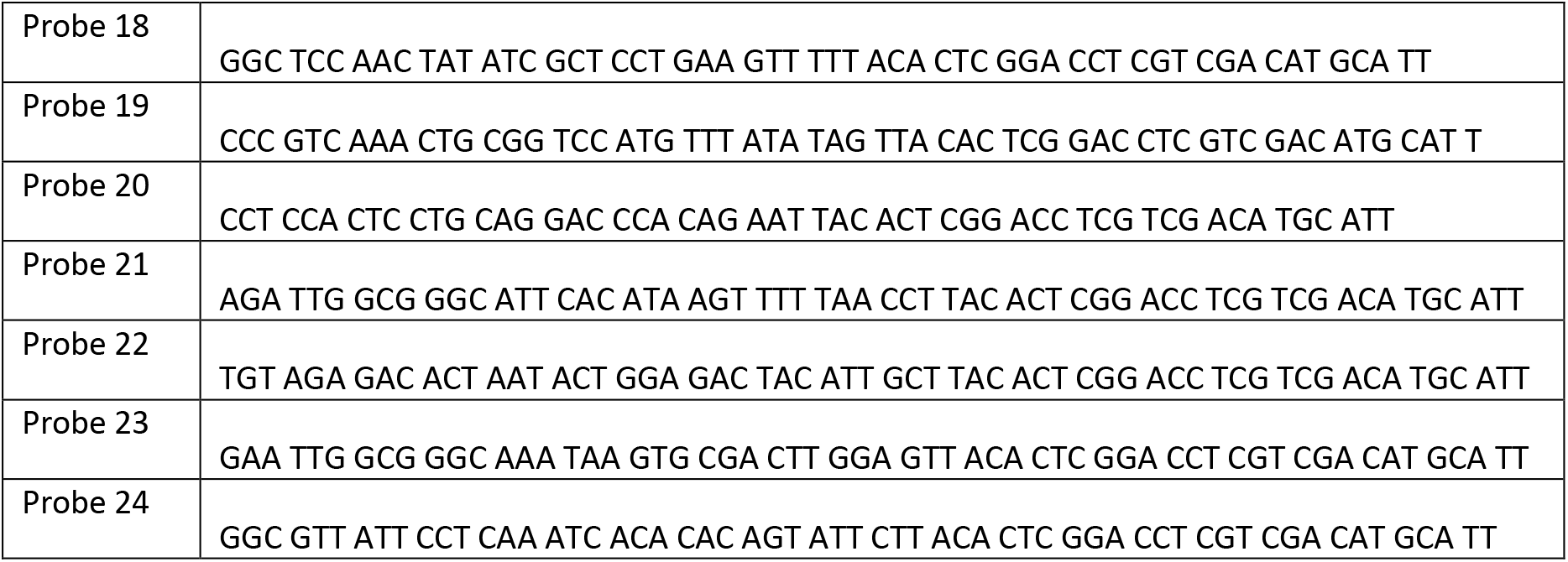

### Fixed Sample Imaging and analysis

Fixed sample imaging was performed on an LSM 880 with Airyscan module (Zeiss). Z planes were acquired with 0.20μm spacing to a typical depth of 80-100 Z-planes from the apical surface of the embryo, using laser scanning confocal in Airyscan super-resolution mode with a zoom of 3.0. DAPI excitation was performed with a 405nm laser and Cy3 excitation with a 561nm laser, with detection on a GaAsP-PMT array coupled to an Airyscan detector. Airyscan processing was performed using 3D Zen Black (Zeiss) prior to analysis. Embryos were staged based on membrane invagination. Z-stacks were taken at both the centre of the presumptive mesoderm as well as at the border region.

Images were analyzed (**Supplemental Figure 3**) with Imaris v9.2.2 by first determining the threshold of detection using the non-mesodermal border region. After applying the threshold to the centre pattern, fixed-sized shells (XY radius >0.3μm, Z radius of >1.0μm) were created around the centroid of detected objects. The median signal intensity of object shells was used as a proxy for the intensity of single molecules of RNA. The transcription site intensity of each nucleus was summed to account for the presence of sister chromatids and treated as a single transcription site throughout. The mean transcription site intensity was divided by the median single molecule intensity to determine the average number of mRNA molecules present at the transcription sites.

### Mathematical modelling of burst parameters and multi-exponential regression fitting

#### Burst Deconvolution and Pol II Positioning

The Pol II positions were found by combining a genetic algorithm with a local optimisation procedure^59^.

Before initiation of the analysis algorithm, several key parameters were established. The Pol II elongation speed was fixed at 45 bp/s^60^. The reporter construct transcript was divided into three sections consisting of the pre-MS2 fragment (41 bp), 24xMS2 loops (1292 bp), and post-MS2 fragment containing the *yellow* reporter (4526 bp). The retention time was assumed to be small in relation to the time needed to produce a transcript, and so was fixed at 0s. The temporal resolution of each movie was 3.86 s/frame. This frame rate is sufficient to detect processes that occur on the order of seconds.

The possible polymerase positions were discretized using a step of 30 bp (or equivalently 2/3 s). This step was chosen, as it is smaller than the minimum polymerase spacing and large enough to have a reasonable computation time. For a movie of 35 min length this choice corresponds to a maximum number of 3150 positions.

The algorithm was implemented in Matlab R2020a using Global Optimization and Parallel Computing Toolboxes for optimizing Pol II positions in parallel for all nuclei in a collection of movies. The resulting positions are stored for analysis in the further steps of our computational pipeline. At this step, the density of Pol II initiation events can be visualised by binning time and checking the occurrence of Pol II activation in each bin. This was rendered as a heatmap in which rows represent a single nucleus time series and the number of activation events per 30 seconds bin (or equivalently 1350 bp) is indicated by the color (**Figure 2E**).

The deconvolution step is common to all of the MS2 data analysis pipelines. A detailed description of the algorithm can be found in^59^.

#### Multi-exponential regression fitting of the survival function and model reverse engineering using the survival function

Data from several movies corresponding to the same genotype was first pooled together. The entry and exit of each trace corresponding to a unique nucleus were defined using a threshold representing 1/5 of the maximum intensity for the specific trace, in order to restrict the analysis to the stable part of the signal (**Figure 2A, grey box**). Waiting times were extracted as differences between successive Pol II positions from all the resulting traces and the corresponding data was used to estimate the nonparametric cumulative distribution function by the Meyer-Kaplan method. This also permits the calculation of a 95% confidence interval for the experimental survival function that is further used to judge the quality of a parametric multi-exponential regression fitting.

Then, a multi-exponential regression fitting produced a set of 2N-1 distribution parameters, where N is the number of exponentials in the regression procedure (3 for N=2 and 5 for N=3). The regression procedure was initiated with multiple initial guesses and followed by local gradient optimisation of the following objective function^59^:

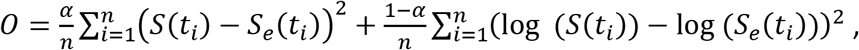

where *S*(*t*_*i*_), *S*_*e*_(*t*_*i*_) are the theoretical (multi-exponential) and empirical (estimated by the Meyer-Kaplan method) survival functions, respectively, and *α* is a parameter satisfying 0 ≤ *α* ≤ 1 and representing the weight of linear scale differences in the objective function. We chose an intermediate value *α* = 0.6 for all our parameter estimates (these estimates are nevertheless robust with respect to *α*).

The optimisation resulted in a best-fit solution with additional suboptimal solutions (local optima with objective function value larger than the best fit). A multi-exponential regression is considered acceptable if the predicted survival function is within the confidence bounds of the experimental survival function. This provides a method to select the number N of exponentials in the regression: we progressively increase N starting with N=2 until an acceptable regression is reached.

The 2N-1 distribution parameters can be computed from the 2N-1 kinetic parameters of a N state transcriptional bursting model. Conversely, a symbolic solution for the inverse problem was obtained, allowing computation of the kinetic parameters from the distribution parameters and reverse engineering of the transcriptional bursting model. In particular, it is possible to know exactly when the inverse problem is well-posed, i.e. there is a unique solution in terms of kinetic parameters for any given distribution parameters in a domain.

The transcriptional bursting models used in this paper are as following (see^59^ for a detailed description):

For N=2, there were 3 distribution parameters and 3 kinetic parameters.

The distribution parameters are *A*_1_, *λ*_1_, *λ*_2_, defining the survival function

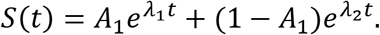

The solution of the inverse problem for the ON-OFF telegraph model (**Figure 2K** and **Figure 6B**) is

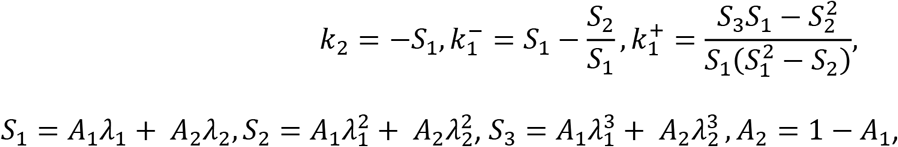

where 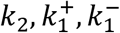 are the initiation rate, the OFF to ON and ON to OFF transition rates, respectively. Thus, the durations of the OFF and ON states can be calculated as:

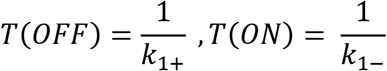

For this model, the probability to be in the state ON and OFF is:

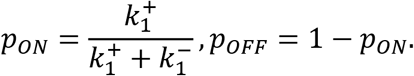

For N=3, there were 5 distribution parameters and 5 kinetic parameters^59^.

The distribution parameters are *A*_1_, *A*_2_, *λ*_1_, *λ*_2_, *λ*_3_, defining the survival function

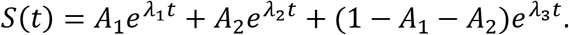

The inverse problem has a unique solution for the 3 state model (non-obligatory pause) with two OFF states (OFF and PAUSE) and one ON state (**Figure 6B’**)

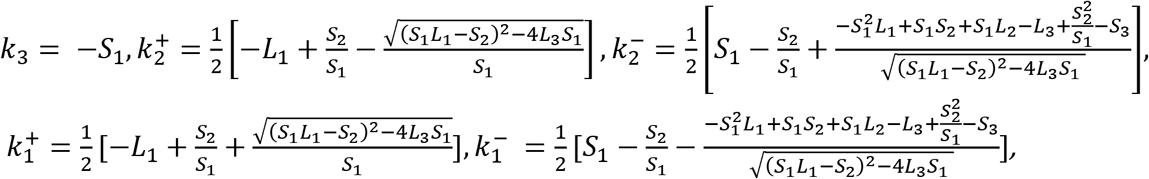

where

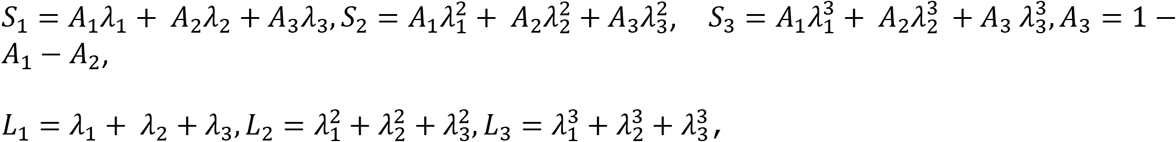

and *k*_3_,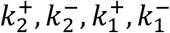 are the transcription initiation, OFF to ON, ON to OFF, PAUSE to ON, and ON to PAUSE rates, respectively.

Duration of the ON, OFF, and PAUSE states can be calculated thusly:

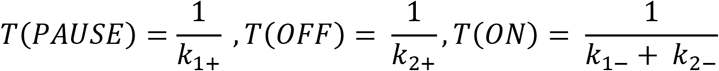

For this model, the steady state probability to be in a given promoter state is

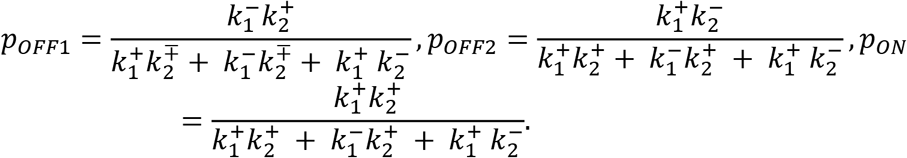

The alternative 3 state model with systematic pause (**Supplemental Figure 5**) satisfies the following relation among distribution parameters^59^:

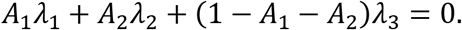

This means that only 4 and not 5 distribution parameters are free, which further constrains the three exponential fitting. In order to infer this model, a constrained fitting was performed but the bad quality of fitting recommended rejection of the model (**Supplemental Figure 5**).

#### Testing the method with artificial data

The entire computational pipeline was tested using artificial data (see also ^59^). Artificial traces were generated by simulating the model using the Gillespie algorithm with parameter sets similar to those identified from data. The simulations generated artificial polymerase positions, from which a first version of the signal was computed by convolution. The results are provided in **Figure 2E-H.**

#### A modified Kolmogorov-Smirnov test for the parametric distribution

A one-sample Kolmogorov-Smirnov (KS) test was used to check if the waiting times follow the parametric fitted distribution. The output of this test is a p-value that is large if the waiting times follow the fitted distribution, and small if not. The KS test is based on differences between estimated and empirical probabilities, being thus sensible to errors in the larger probabilities of shorter waiting times, and less sensitive to rare, very long waiting times. Our fitting procedure combines linear and logarithmic scales to find a balance between short and long time scales (see ^59^). Although relatively small (in terms of the objective function), the resulting errors on short times are large enough to be considered significant by the KS test for all models. In order to be able to distinguish between models, we have limited the analysis to times larger than 10-20s.

#### Error intervals

Distribution parameters result from multi-exponential regression fitting using gradient methods with multiple initial data. These optimization methods provide a best fit (global optimum) but also suboptimal parameter values. Using an overflow ratio (a number larger than one, in our case 2) to restrict the number of suboptimal solutions, we define boundaries of the error interval as the minimum and maximum parameter value compatible with an objective function less than the best fit times the overflow.

### Mathematical modelling of post-mitotic gaps

The distribution of the post-mitotic gap was estimated using the equation below. The fitting suggests that the timescale parameters of the post-mitotic gap are even, i.e. *λ*_1_ = *λ*_2_ = *λ*_3_. In this limit, the five parameters, three exponential distribution defined by the equation below, becomes the simpler, three parameters mixed gamma distribution described as

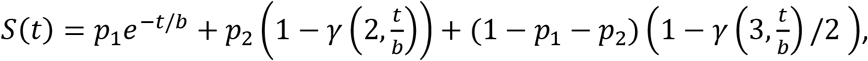

where 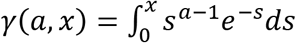 is the lower incomplete gamma function, and *p*_1_, *p*_2_, *b*, are the probabilities of one step, two steps, and the mean step duration, respectively.

### Chromatin Immunoprecipitation

Homozygous embryos were collected and fixed in 1.4% formaldehyde for 20 minutes prior to dry storage at −80°C. Embryos were dissociated on ice in RIPA buffer (150mM NaCl, 1.0% IGEPAL CA-630, 0.5% sodium deoxycholate, 0.1% SDS, 50mM Tris-HCl pH 7.8) supplemented with protease and phosphatase inhibitors (Roche). Chromatin shearing was performed in a pre-chilled Bioruptor Pico (Diagenode) for 5 cycles of 30s/30s. Samples were divided into equal volumes and incubated overnight at 4°C with either rabbit anti-Rbp3 or rabbit anti-Nelf-E (gifts from J. Zeitlinger ^16^) or normal serum. A second incubation overnight at 4°C with Protein G-magnetic Sepharose beads (GE Healthcare) was used to pull down protein:DNA complexes prior to washing in low salt (120mM NaCl, 0.1% SDS, 0.5% Triton X-100, 20mM Tris-HCl pH 8.0) and high salt (500mM NaCl, 0.1% SDS, 0.8% Triton X-100, 20mM Tris-HCl pH 8.0) buffers, elution, incubation with protease K and RNAse A, and DNA retrieval using the QiaQuick PCR cleanup kit (Qiagen). qPCR analysis was performed using Light Cycler 480 SYBR Green I Master Mix (Roche). Analysis was performed using Microsoft Excel and Prism (Graphpad) with a Mann-Whitney U test to determine significance.

To test RNAi-mediated knockdown of Nelf-A expression, 0-2h embryos were homogenized in Trizol (Invitrogen) and RNA was extracted per manufacturer’s directions. Reverse transcription was performed using the Superscript IV system (Invitrogen) with oligo d(T)20 priming prior to qPCR analysis. *nos:*GAL4-VP16; UAS:*white* RNAi embryos were used as the control and all measurements were performed in biological duplicate and technical triplicate. qPCR analysis was performed using Light Cycler 480 SYBR Green I Master Mix (Roche). Analysis was performed using Microsoft Excel and Prism (Graphpad) with a Student’s T-test to determine significance.

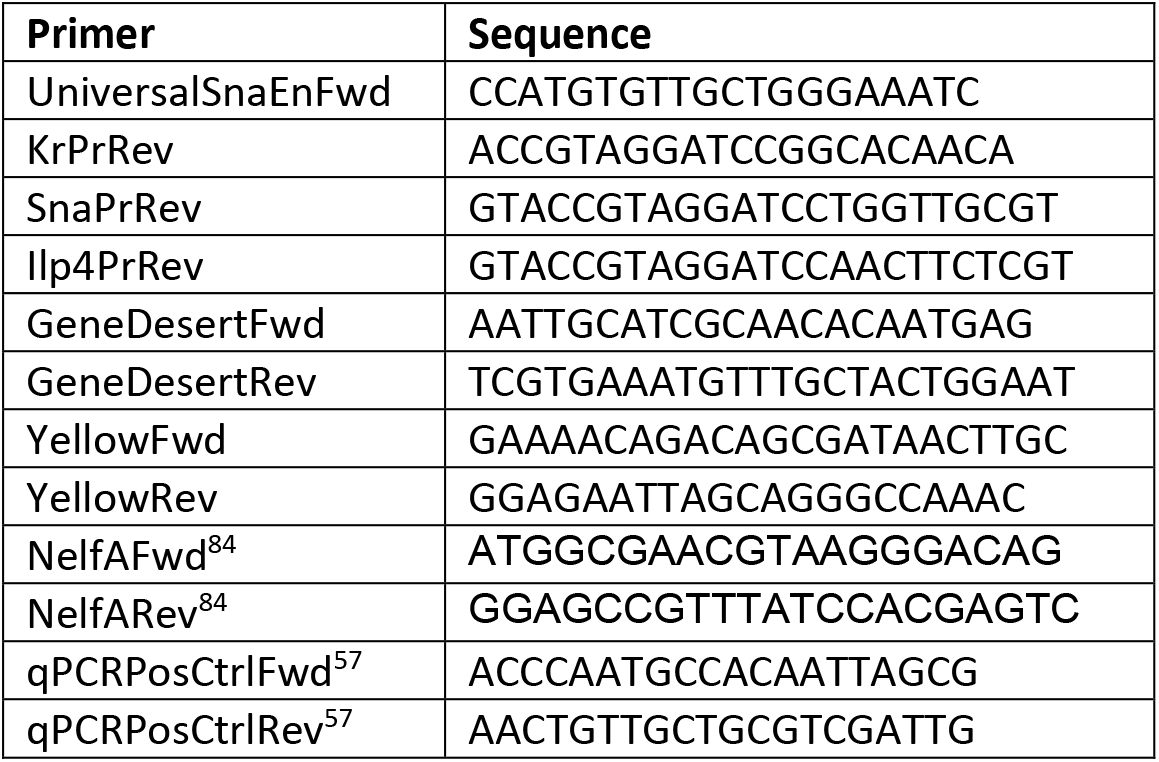

### Software/Code Availability

Live imaging analysis software is available at http://www.igmm.cnrs.fr/segment-track/ along with a video tutorial. Code for the mathematical analysis of burst parameters and multi-exponential regression fitting is available^59^.

---

**Supplemental Figure 1:**
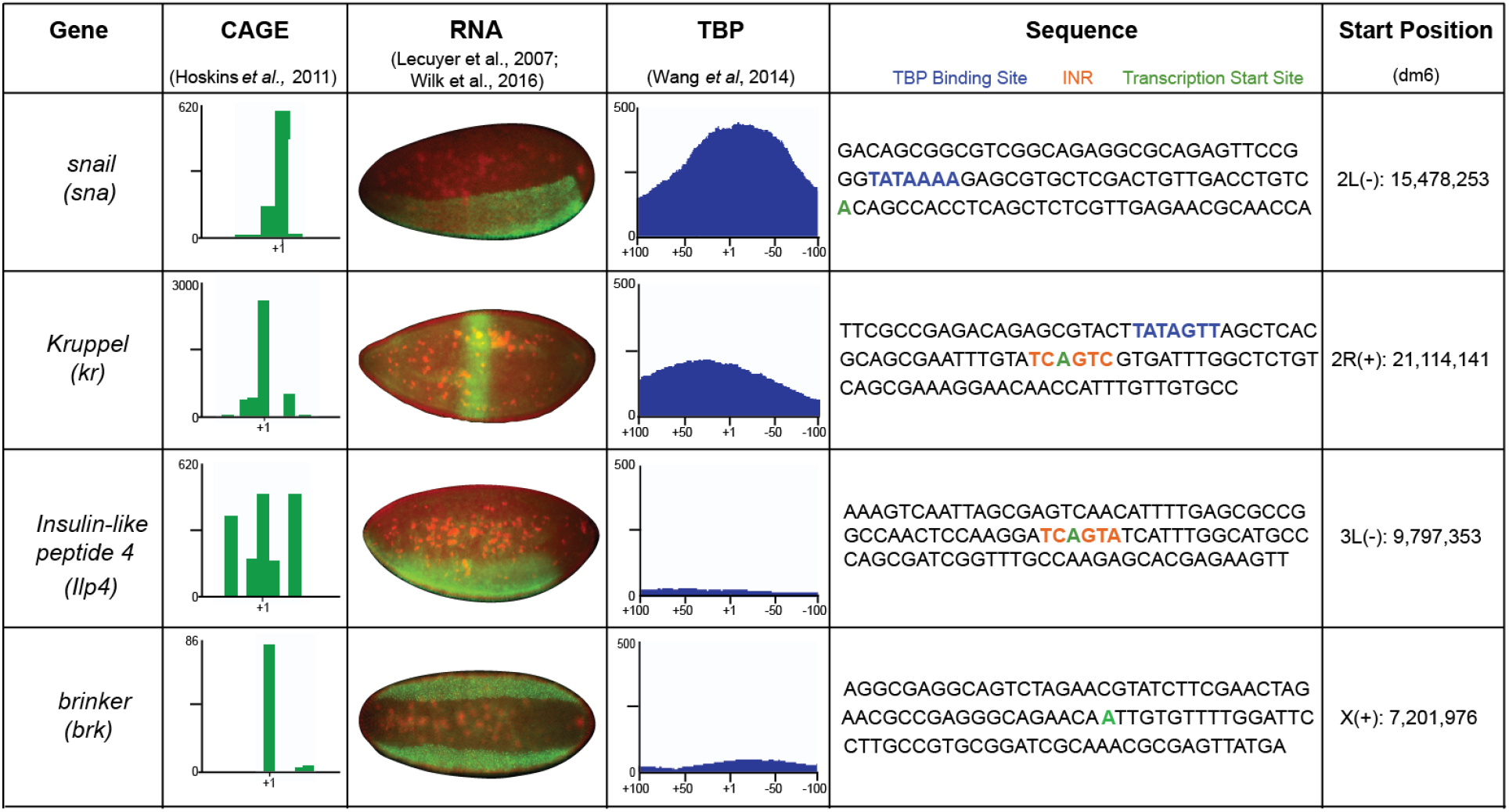
Summary of *sna*, *kr*, *Ilp4* and *brk* promoters. CAGE profile^54^, representative nc14 mRNA expression pattern^52,53^, TBP binding profile^85^, annotated promoter sequence and genomic coordinates for the developmental promoters used in this study. For each of the four developmental promoters, the highest peak in the CAGE profile indicates the most used transcription initiation site.

**Supplemental Figure 2:**
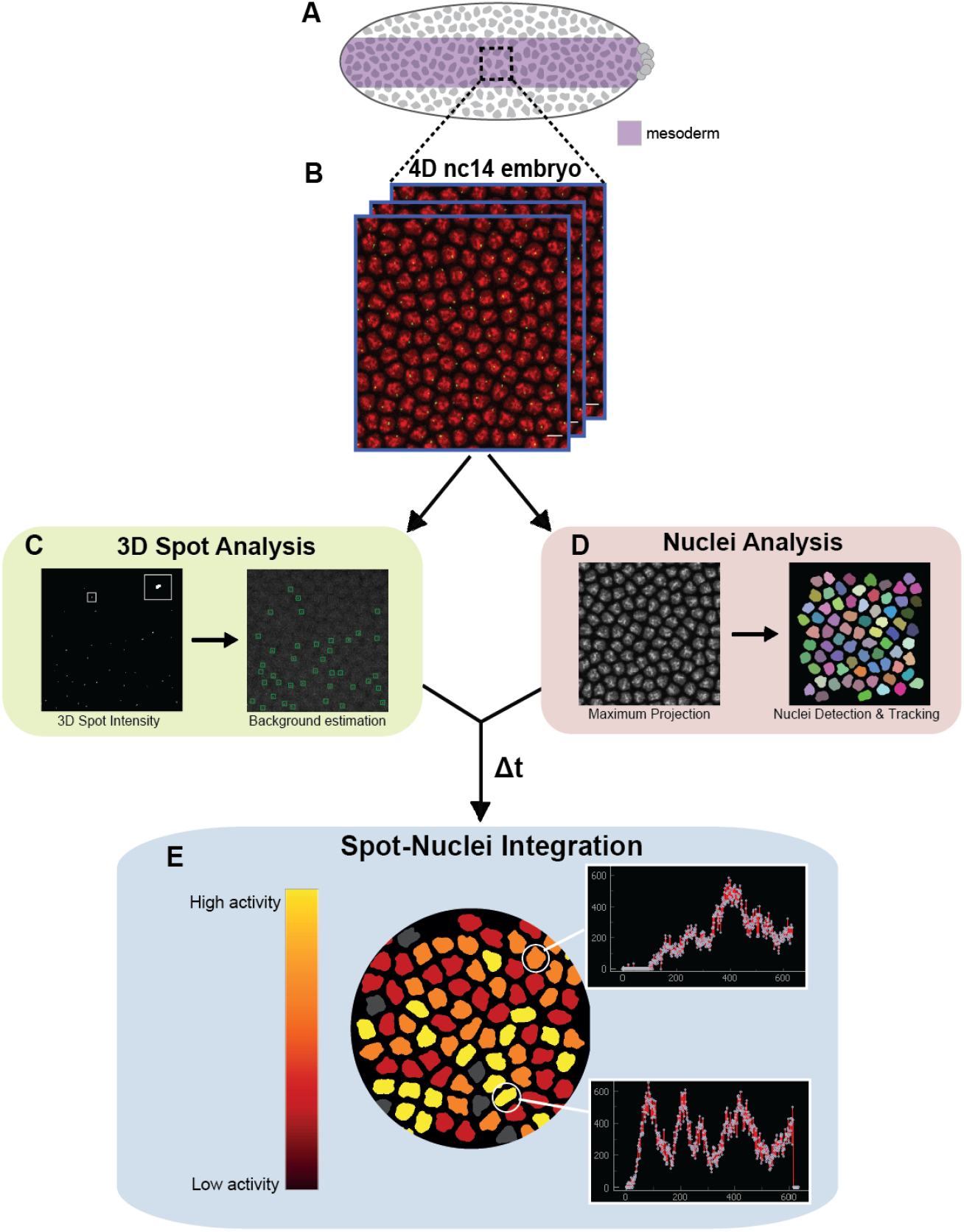
Image analysis pipeline for retrieving single nuclei transcriptional activities. **A)** Schematic of the imaging set-up indicating spatial restriction to presumptive mesoderm (purple). **B)** Representation of Z-series showing nuclei (red) and MS2/MCP-GFP puncta (green). Scale bar is 5μm. **C)** Spot analysis is performed in 3D for each time point in the Z series. Inset shows zoom on single detected spot in software interface. Background estimation is performed independently for each spot using an average intensity value of the surrounding pixels (green rectangle). Nuclei analysis is performed in 2D for each time point. Nuclei are maximum projected, smoothed and detected, and then tracked between time points. Nuclei at the border are removed. **E)** Spot and nuclei information are integrated to associate each nuclei with its nearest transcription site for each time point in the Z-series, which can be mapped to each nuclei within the pattern.

**Supplemental Figure 3:**
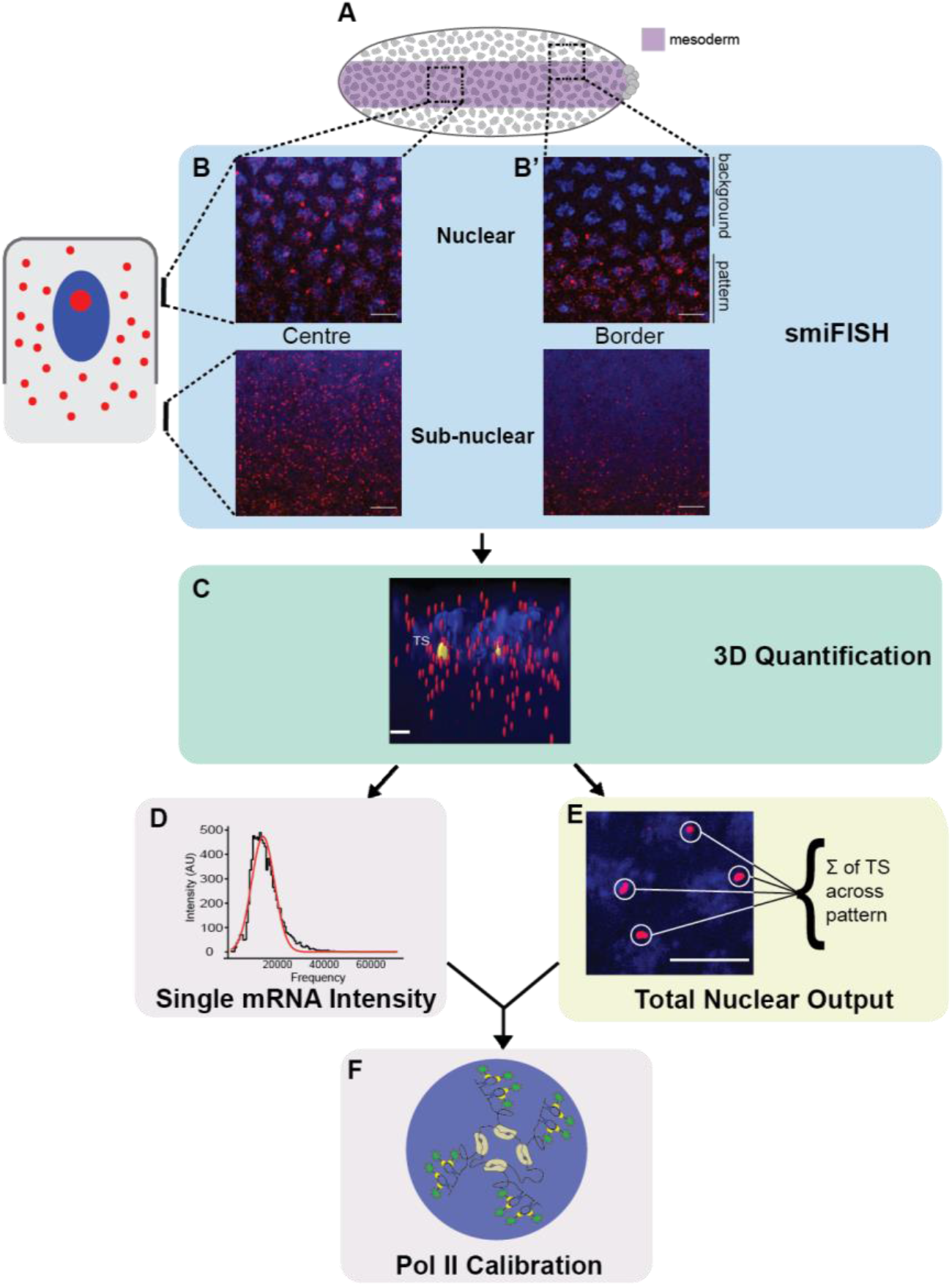
Image analysis pipeline for single molecule FISH. **A)** Schematic of the embryo showing center and border imaging regions at the presumptive mesoderm (purple). **B)** Single molecule inexpensive fluorescence *in situ* hybridization (smiFISH)^83^ was used to identify individual mRNAs as well as the transcription site at the nuclei. Images are of a 1μm Z stack centred on the nuclear and subnuclear regions with DAPI in blue and mRNA in red. Scale bar is 5μm. **C)** 3D quantification of single molecule and transcription site intensity with Imaris software (v9.2.2). Intensities of single molecules (red) and transcription sites (TS, yellow) around the nuclei (blue) were independently measured. **D) E**stablishment of the median single molecule intensity for the entire Z-stack. **E)** Transcription site intensities across the pattern were summed and divided by the number of active nuclei such that sister chromatids were resolved as a single site, and the average TS intensity was established^86^. **F)** The mean TS intensity of a Z-stack was divided by the median single molecule intensity of the Z stack to give an average number of transcripts per transcription site. At least 3 independent replicates per genotype were analyzed.

**Supplemental Figure 4:**
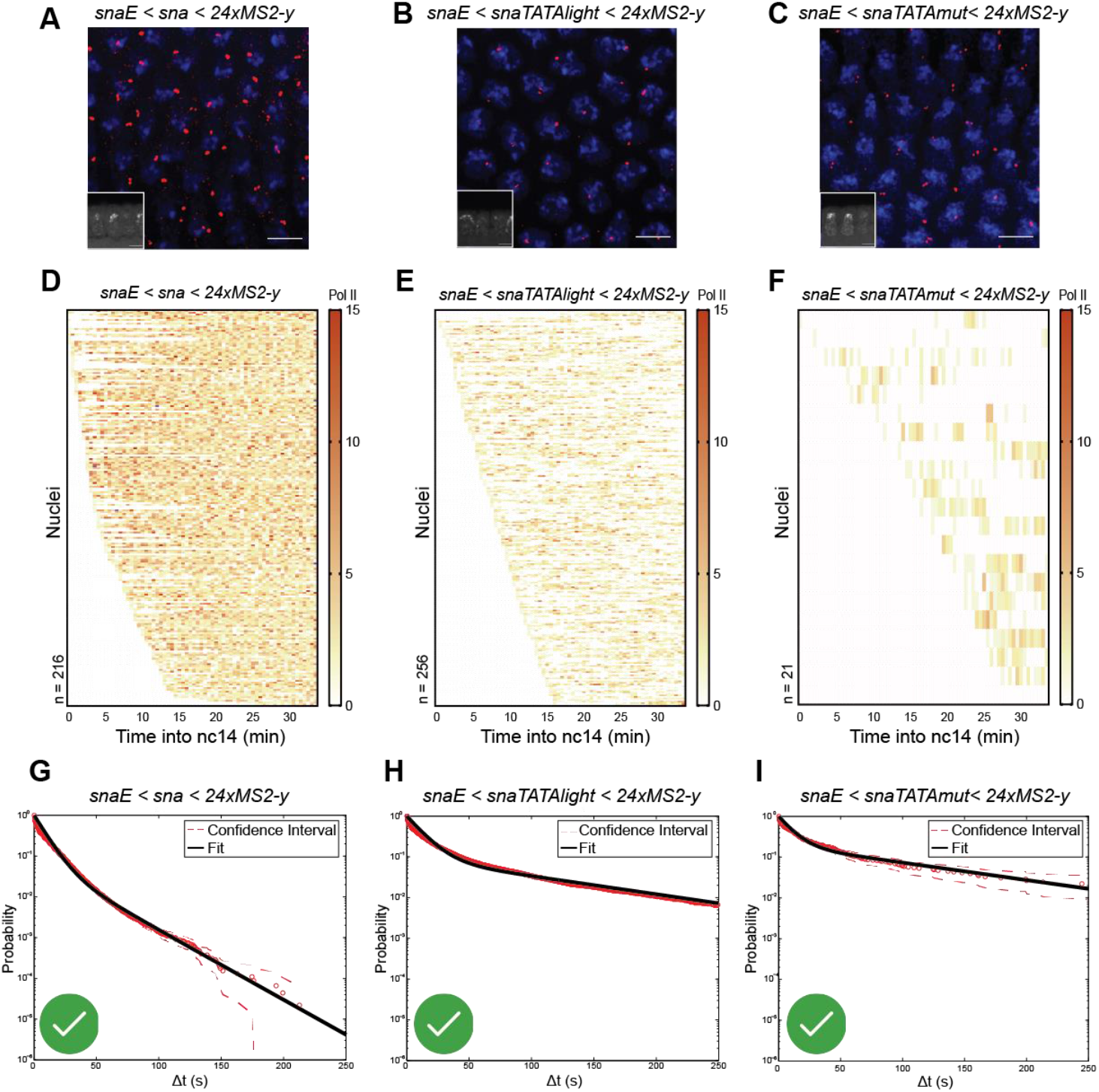
Impact of the TATA box on *sna* promoter dynamics. **A-C)** Nuclear cycle 14 embryos, expressing one copy of the *snaE<sna* (A), *sna<snaTATAlight* (B) or the *sna<snaTATAmut* (C) transgene stained with *yellow* probes (smiFISH) and DAPI. Images are representative confocal 1μm z-stack at the plane of the nuclei to show transcription site foci. Note that in some nuclei, two spots presumably emanating from two apposed sister chromatids are observable. Insets in A, B and C: zoomed view showing membrane invagination. Scale bar is 5 μm in all images. **D-F)** Heatmap showing frequency of Pol II initiation events per 30s window during the first 30 minutes of nc14 for each of the genotypes indicated. **G-I)** Survival function of the distribution of waiting times between polymerase initiation events (red circles) and the two-exponential fitting of the population (black line) for *sna* (p-value KS test = 0.717), *snaTATAlight* (p = 5.99e^−4^) and *snaTATAmut* (p-value KS test = 0.820). Green check indicates accepted fitting. **Statistics:** *snaE<sna*, 216 nuclei, 3 movies; *snaE<snaTATAlight*, 256 nuclei, 6 movies; *snaE<snaTATAmut*, 21 nuclei, 3 movies. See **Supplemental Movies S1-S3 and Supplemental Table 2)**.

**Supplemental Figure 5:**
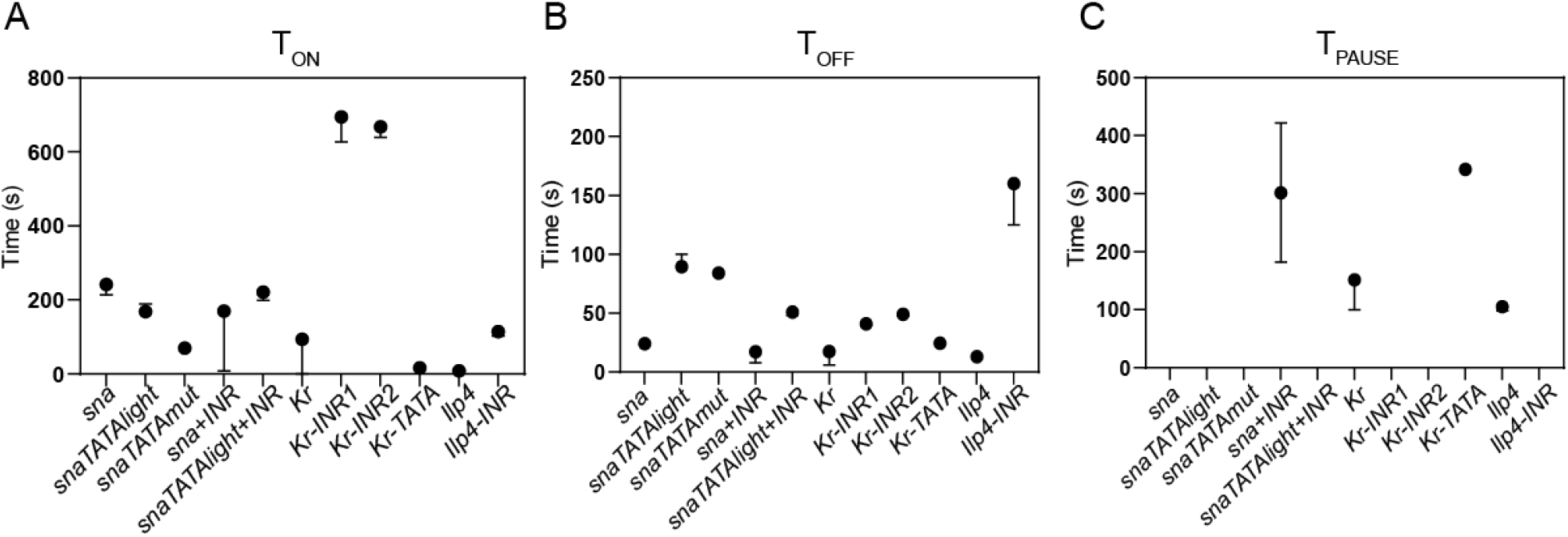
Average State Durations and Probabilities. **A)** Average duration of the ON state of best fitting for indicated promoters. Error bars represent the error interval. **B)** Average duration of the OFF state of best fitting for indicated promoters. Error bars represent the error interval. **C)** Average duration of the PAUSE state of best fitting for indicated promoters. Error bars represent the error interval.

**Supplemental Figure 6:**
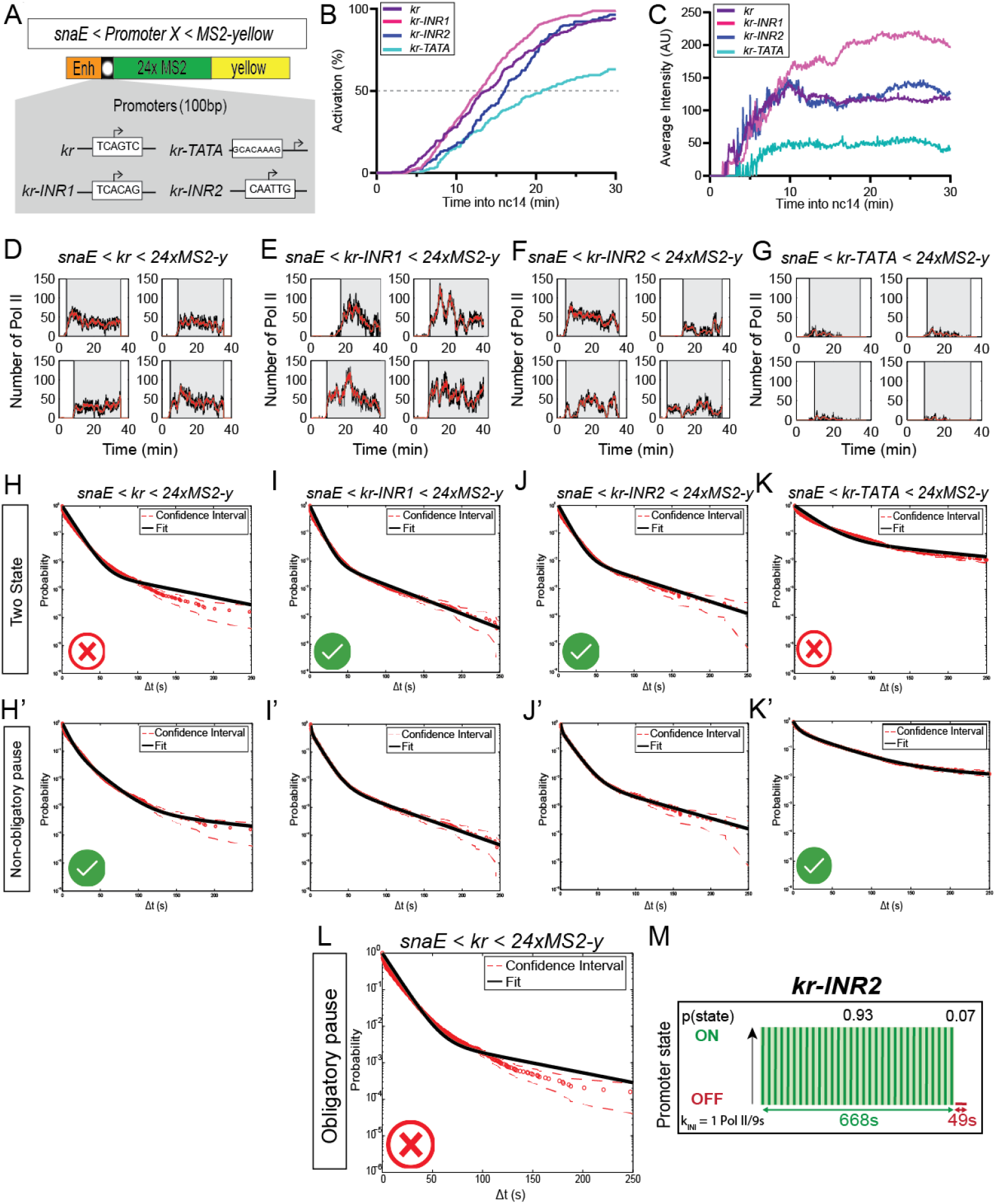
Elucidating the impact of INR for *kr* promoter dynamics. **A)** Schematic of the *kr* promoter transgenic variants. Two mutants of the INR motif were generated, *kr-INR1* corresponds to *kr* INR swap with the TSS region of *sna*, whereas *kr-INR2* is the swap with the TSS region of *brk*. *kr-TATA* contains the same TBP binding site mutation as the *snaTATAmut* promoter upstream of the indicated TSS. **B)** Cumulative activation curves of all nuclei during the first 30 minutes of nc14 for indicated genotypes. Time zero is from anaphase during nc13-nc14 mitosis. **C)** Average instantaneous fluorescence of transcriptional foci of active nuclei during the first 30 minutes of nc14 for indicated genotypes. Time zero is from anaphase during nc13-nc14 mitosis. **D-G)** Examples of individual activity events for the indicated genotype. Red lines represent the reconstructed signal after Pol II positioning. **H-K)** Survival function of the distribution of waiting times between polymerase initiation events (red circles) and the two-exponential (H-K) and three exponential (H’-K’) fitting of the population (black line). **K, cont’d)** Red dashes represent 95% confidence interval. p-values of the KS test for two state fittings: *kr* p-value = 2.1e^−5^, *kr-INR1* p-value = 0.941, *kr-INR2*, p-value = 0. 980, *kr-TATA* p-value = 4.21e^−16^. p-values of the KS test for three state fittings: *kr* p-value = 0.508, *kr-INR1* p-value = 0.999, *kr-INR2*, p-value = 0. 999, *kr-TATA* p-value = 0.978 (see also **Supplemental Table 1**). Green check indicates accepted fitting, red X indicates rejected fitting. **L)** Survival function of the distribution of waiting times between polymerase initiation events (red circles) and the three exponential obligatory pause model fitting of the population (black line). Red dashes represent 95% confidence interval. **M)** Representation of the *kr-INR2* promoter dynamics. ON state durations are depicted in green and OFF states in red with state probabilities shown above. **Statistics:** *snaE<kr*, 243 nuclei, 3 movies; *snaE<kr-INR1*, 342 nuclei, 4 movies; *snaE<kr-INR2*, 168 nuclei, 3 movies; *snaE<kr-TATA*, 98 nuclei, 2 movies. See **Supplemental Movie S5-S6** and **Supplemental Table 2**.

**Supplemental Figure 7:**
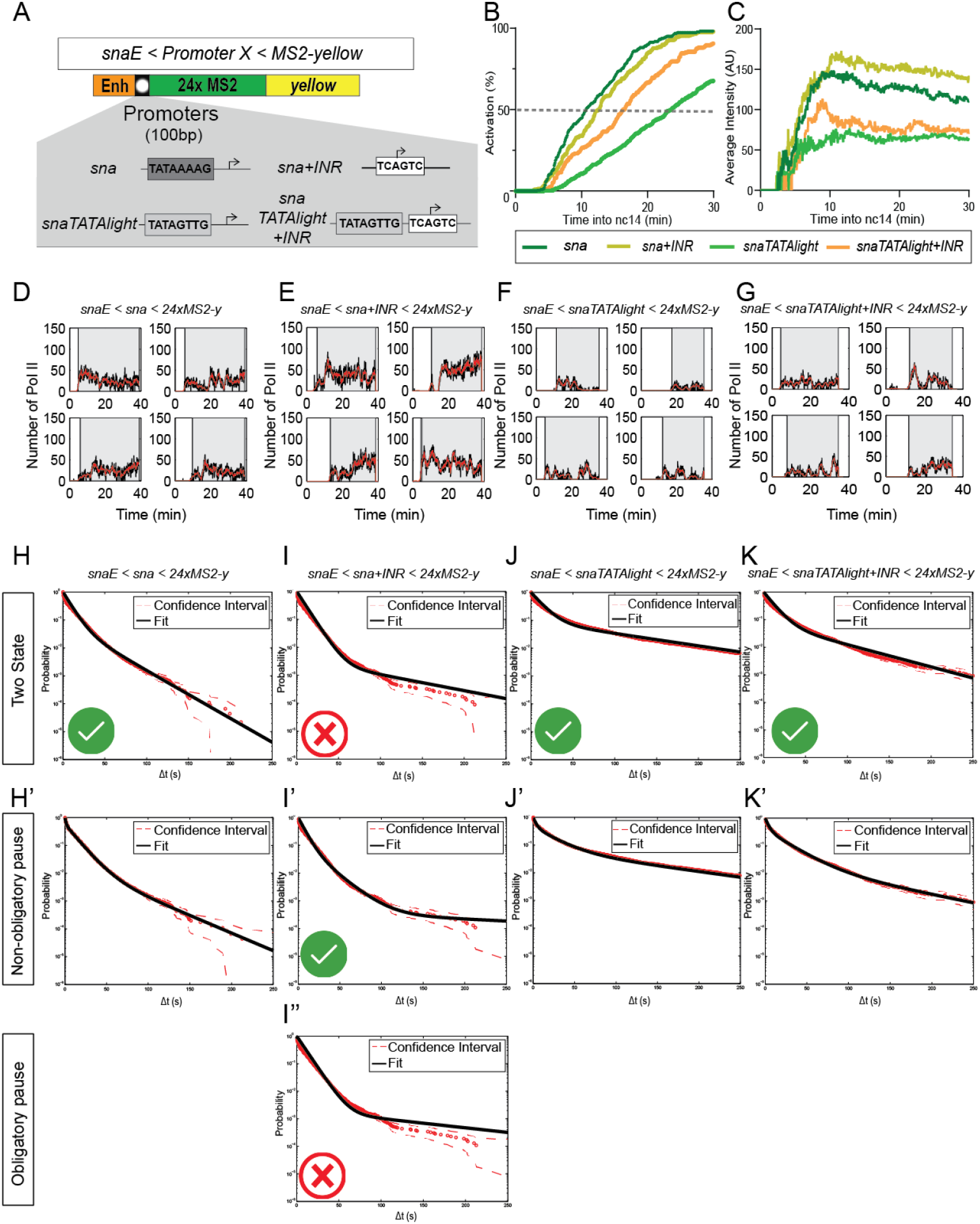
Impact of the INR and TATA box on *sna* promoter dynamics. **A)** Scheme of the *sna* promoter and the 3 mutants generated to decode the role of the TATA-box and the INR motif: *snaTATAlight*, *sna+INR*, *snaTATAlight+INR*. **B)** Cumulative activation curves of all nuclei during the first 30 minutes of nc14 for indicated genotypes. Time zero is from anaphase during nc13-nc14 mitosis. **C)** Average instantaneous fluorescence of transcriptional foci of active nuclei during the first 30 minutes of nc14 for indicated genotypes. Time zero is from anaphase during nc13-nc14 mitosis. **D-G)** Examples of single nuclei traces for the indicated genotypes. Red lines represent the signal after Pol II positioning. **H-K)** Survival function of the distribution of waiting times between polymerase initiation events (red circles) and the two-exponential (H-K), three exponential non-obligatory paused (H’-K’) and three exponential obligatory pause (I”) fitting of the population (black line). Red dashes represent 95% confidence interval. p-values of the KS test for two state fittings: *sna* p = 0.717, *sna+INR* p = 1.37e^−4^, *snaTATAlight* p = 5.99e^−4^, *snaTATAlight+INR* p-value = 8.39e^−2^. p-values of the KS test for three state non-obligatory fittings: *sna* p-value = 0.925, *sna+INR* p-value = 0.999, *snaTATAlight* p-value = 0.370, *snaTATAlight+INR* p-value = 0.997 (see **H-K, cont’d)** also **Supplemental Table 1**). Green check indicates accepted fitting, red X indicates rejected fitting. **Statistics:** *snaE<sna*, 216 nuclei, 3 movies; *snaE<sna+INR*, 236 nuclei, 4 movies; *snaE<snaTATAlight*, 353 nuclei, 6 movies; *snaE<snaTATAlight+INR*, 193 nuclei, 3 movies. See **Supplemental Movies S1-S2, S4 and Supplemental Table 2**.

**Supplemental Figure 8:**
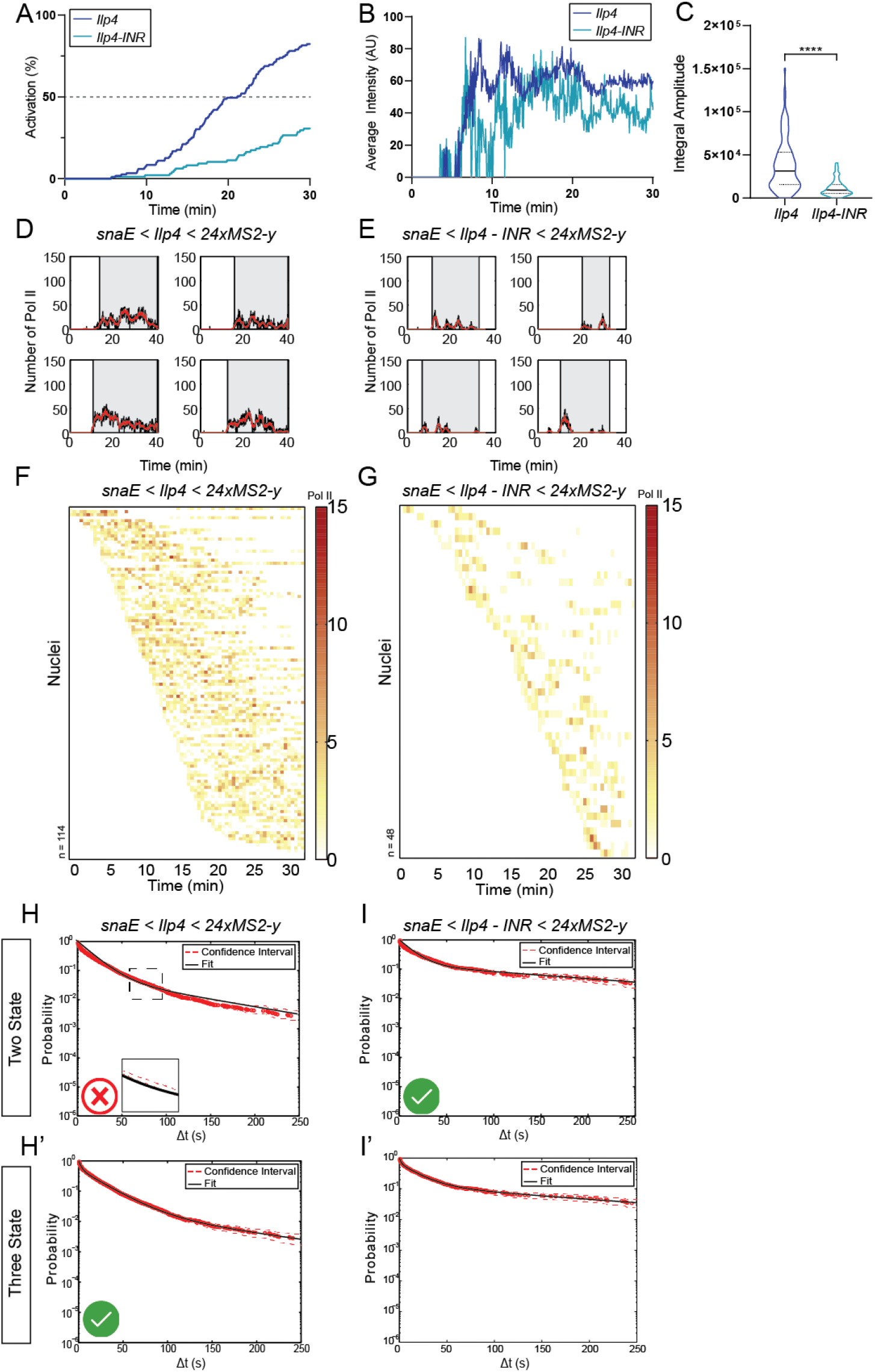
Impact of the INR on *Ilp4* promoter dynamics. **A)** Cumulative activation curves of all nuclei during the first 30 minutes of nc14 for indicated genotypes. Time zero is from anaphase during nc13-nc14 mitosis. **B)** Average instantaneous fluorescence of transcriptional foci of active nuclei during the first 30 minutes of nc14 for indicated genotypes. Time zero is from anaphase during nc13-nc14 mitosis. **C)** Distribution of individual trace integral amplitudes from first 30 minutes of nc14. Solid line represents median and dashed lines the first and third quartiles, using a Kruskal-Wallis test for significance with multiple comparison adjustment. **** p < 0.0001 **D-E)** Examples of individual activity events for the indicated genotypes. Red lines represent the reconstructed signal after Pol II positioning. **F-G)** Heatmaps indicating number of polymerases during nc14 for each genotype. **H-I)** Survival function of the distribution of waiting times between polymerase initiation **H-I, cont’d)** events (red circles) and the two-exponential (H-I) and three exponential (H’-I’) fitting of the population (black line). Red dashes represent 95% confidence interval. p-values of KS test for two state fittings: *Ilp4* p-value = 1.42e^−08^, *ilp4-INR* p-value = 0.902. p-values of KS test for three state fittings: *Ilp4* p-value = 0.998, *ilp4-INR* p-value = 0.999 (see also **Supplemental Table 1**). Green check indicates accepted fitting, red X indicates rejected fitting. **Statistics:** *snaE<Ilp4*, 114 nuclei, 2 movies; *snaE<Ilp4-INR*, 48 nuclei, 2 movies. See **Supplemental Table 2**.

**Supplemental Figure 9:**
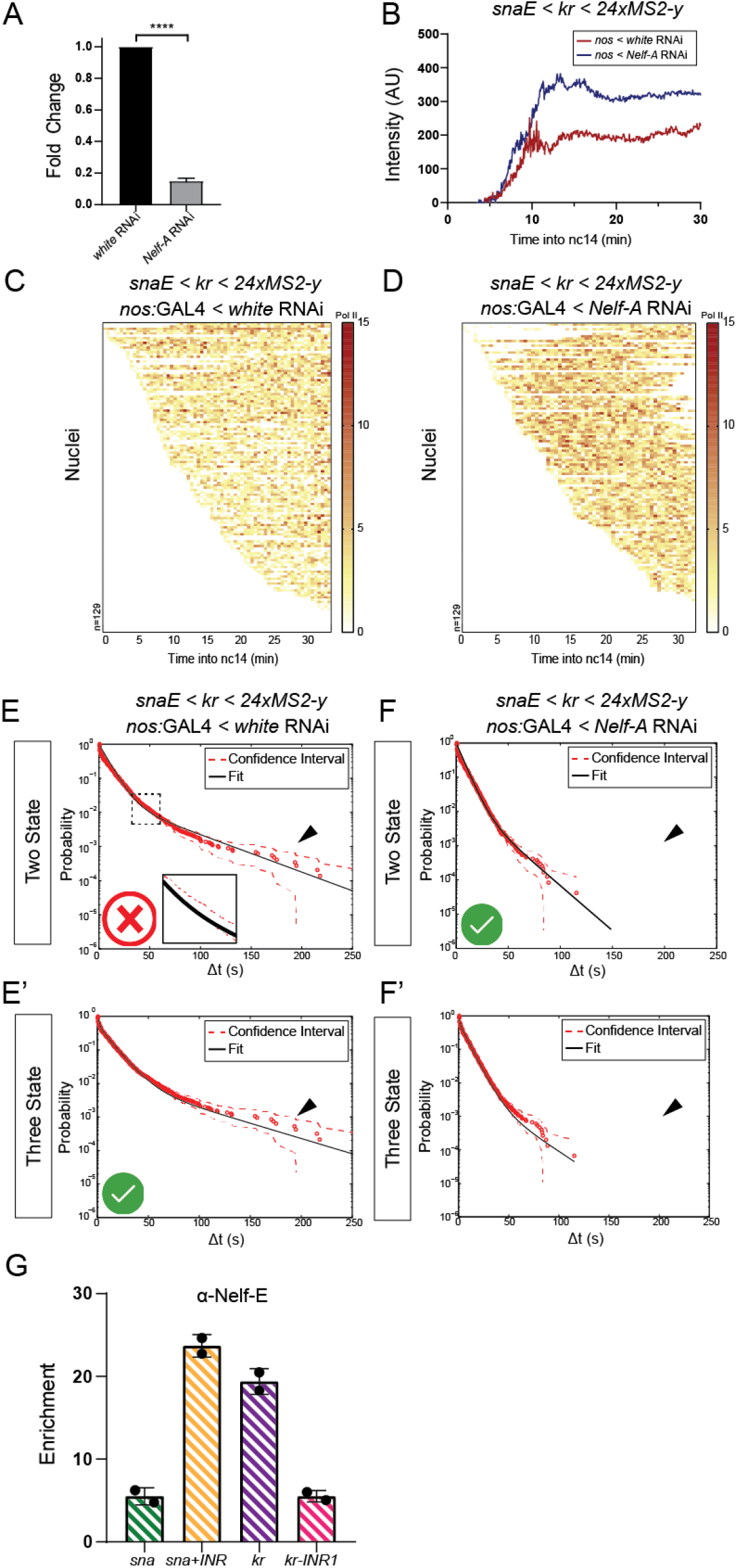
Altering pausing in *trans* changes state dynamics. **A)** RNAi-mediated knockdown of *Nelf-A* transcript level measured by qRT-PCR (average ± SD). n=3 biological replicates, **** p<0.0001 using a Student’s t-test. **B)** Average instantaneous fluorescence of transcriptional foci of active nuclei during the first 30 minutes of nc14 for indicated genotypes. Time zero is from anaphase during nc13-nc14 mitosis. **C-D)** Heatmaps indicating number of polymerases during nc14 for each genotype. **E-F)** Survival function of the distribution of waiting times between polymerase initiation events (red circles) and the two-exponential (E-F) and three exponential (E’-F’) fitting of the population (black line). Red dashes represent 95% confidence interval. p-value of the KS test for two state fittings: *white* RNAi; *kr* p-value = 5.30e^−14^, *Nelf-A* RNAi*; kr* p-value = 2.51e^−2^. p-values of the KS test for three state fittings: *white* RNAi; *kr* p-value = 0.237, *Nelf-A* RNAi*; kr* p-value = 0.674 (see also **Supplemental Table 1**). Green X indicates accepted fitting, red X indicates rejected fitting. Arrowheads indicate long duration waiting times. **G)** Enrichment of Nelf-E at promoter region of indicated transgenes using ChIP-qPCR, shown as average ± SD of n=2 biological replicates with individual replicates indicated. **Statistics:** *nos:GAL4 < white* RNAi; *snaE<kr*, 129 nuclei, 2 movies; *nos:GAL4 < Nelf-A* RNAi; *snaE<kr,* 129 nuclei, 2 movies. See **Supplemental Movies S7-S8**.

**Supplemental Figure 10:**
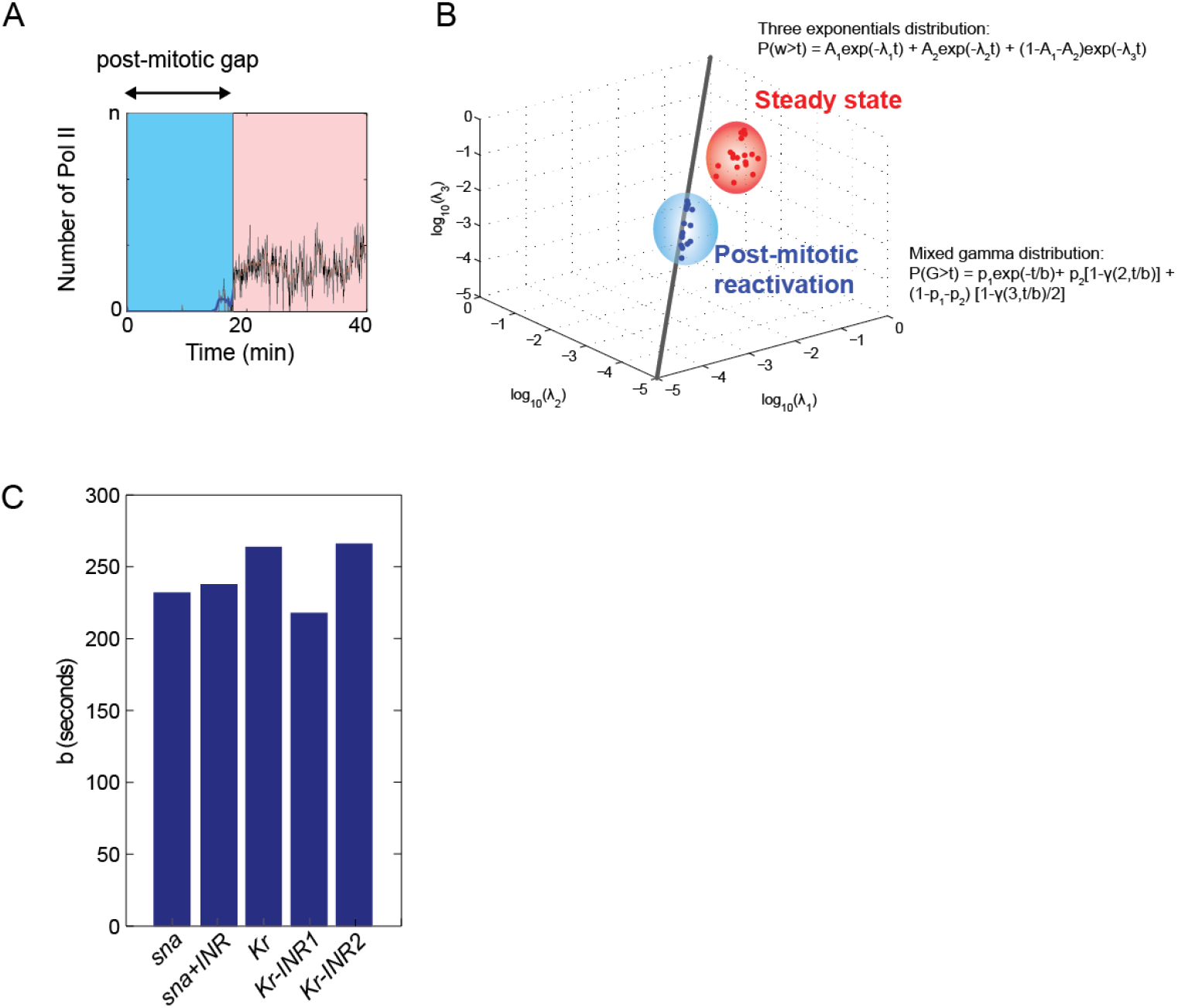
Distribution of the post-mitotic gaps. **A)** Example trace indicating timeframe of post-mitotic gap (G, blue), i.e. the lag time between nc13 to nc14 mitosis to transcriptional activation in nc14. In comparison, the time window used to decipher bursting kinetics with the deconvolution method is shown in red. **B)** A three-exponential fit of the data from various promoters shows that the post-mitotic gaps have different timescales (inverses of λi) than the bursting waiting time. The timescales of the post-mitotic gap duration are even (λ_1_ = λ_2_ = λ_3_ =1/b). Thus, a mixed gamma distribution fits well the distribution of the post-mitotic gap. **C)** The values of the gap step durations (b) for different promoters.

**Supplemental Table 1:** Objective functions and Kolmogorov-Smirnov test results for promoters derived from deconvolution and multi-exponential regression fitting of live imaging data. Parameters are provided for the most parsimonious fitting of each genotype. Bold indicates the most parsimonious appropriate fitting of the data. A two-state model was considered rejected when the p-value of the Kolmogorov-Smirnov test was less than 5.00e-4. *As the confidence interval criterion rejected and the KS criterion accepted the two-state model for the *SnaTATAlight+INR* promoter, this genotype was considered borderline.

**Supplemental Table 2:** Kinetic parameters for promoters derived from deconvolution and multi-exponential regression fitting of live imaging data. Parameters are provided for the most parsimonious fitting of each genotype, with the optimal fitting highlighted in grey. Minimum and maximum values indicate the boundaries of the error interval. State durations are calculated from the provided switching rates (*k*_*n*_) and time durations for each state are provided as ‘T(*state*)’. State probability values are indicated as ‘p(*state*)’. All calculations are provided in **Methods**.

**Supplemental Table 3:** Promoter sequences for the genotypes listed in this study.

**Supplemental Movie S1:** Representative false-colour imaging of *snaE<sna<24xMS2-yellow* in nc14. Nuclei colour corresponds to GFP fluorescence intensity as in **Fig 3C** with inactive nuclei in grey. Intensity scale is common to Supplemental Movies S1-S3.

**Supplemental Movie S2:** Representative false colour imaging of *snaE<snaTATAlight<24xMS2-yellow* in nc14. Nuclei colour corresponds to GFP fluorescence intensity as in **Fig 3C** with inactive nuclei in grey. Intensity scale is common to Supplemental Movies S1-S3.

**Supplemental Movie S3:** Representative false colour imaging of *snaE<snaTATAmut<24xMS2-yellow* in nc14. Nuclei colour corresponds to GFP fluorescence intensity as in **Fig 3C** with inactive nuclei in grey. Intensity scale is common to Supplemental Movies S1-S3.

**Supplemental Movie S4:** Live imaging of *snaE<sna+INR<24xMS2-yellow* representative of NC14 beginning at mitosis. Nuclei are detected using His2Av-mRFP and MS2 using MCP-GFP.

**Supplemental Movie S5:** Live imaging of *snaE<kr<24xMS2-yellow* representative of NC14 beginning at mitosis. Nuclei are detected using His2Av-mRFP and MS2 using MCP-GFP.

**Supplemental Movie S6:** Live imaging of *snaE<kr-INR1<24xMS2-yellow* representative of NC14 beginning at mitosis. Nuclei are detected using His2Av-mRFP and MS2 using MCP-GFP.

**Supplemental Movie S7:** Live imaging of *snaE<kr<24xMS2-yellow* in a *nanos:GAL4> UAS:MCP-GFP-His2Av-RFP, UAS:RNAi white* background, representative of nc14 beginning at mitosis. Nuclei are detected using His2Av-mRFP and MS2 using MCP-GFP.

**Supplemental Movie S8:** Live imaging of *snaE<kr<24xMS2-yellow* in a *nanos:GAL4> UAS:MCP-GFP-His2Av-RFP, UAS:RNAi Nelf-A* background, representative of nc14 beginning at mitosis. Nuclei are detected using His2Av-mRFP and MS2 using MCP-GFP.

